# Lymph Node Fibroblast Phenotypes and Immune Crosstalk Regulated by Podoplanin Activity

**DOI:** 10.1101/2022.12.01.518753

**Authors:** Spyridon Makris, Yukti Hari-Gupta, Jesús A. Cantoral-Rebordinos, Victor G. Martinez, Harry L. Horsnell, Agnesska C. Benjamin, Isabella Cinti, Martina Jovancheva, Daniel Shewring, Nicola Nguyen, Charlotte M. de Winde, Alice Denton, Sophie E. Acton

**Affiliations:** Stromal Immunology Group, MRC Laboratory for Molecular Cell Biology, University College London, Gower Street, London WC1E 6BT, UK; Stem Cell and Human Development Laboratory, The Francis Crick Institute, 1 Midland Road, NW1 1AT; Molecular Oncology Unit, Centro de Investigaciones Energéticas, Medioambientales y Tecnológicas (CIEMAT), Madrid, Spain; Department of Immunology and Inflammation, Imperial College London, London, W12 0NN, UK; Department for Molecular Cell Biology and Immunology, Amsterdam UMC, location VUmc, De Boelelaan 1108, 1081 HZ Amsterdam, Netherlands

**Keywords:** Lymphoid Tissue, Fibroblast, Podoplanin, Inflammation, Stromal immunology

## Abstract

Lymph nodes are uniquely organised niches for immune interactions. Fibroblastic reticular cells (FRCs) facilitate immune cell communication and regulate immune function by producing growth factors, chemokines and inflammatory cues. Stromal expression of the glycoprotein Podoplanin (PDPN) is required for lymph node development, but the requirement for PDPN signaling in FRCs in adult lymph nodes has not been directly tested. Using a conditional *in vivo* deletion model PDGFRα^mGFPΔPDPN^, scRNA-seq revealed that PDPN deletion increased inflammatory signalling in lymph nodes and disrupted stromal-immune cells crosstalk in steady state and during adaptive immune responses. We confirmed that PDPN/CLEC-2 signaling in FRCs switches transcriptional states and alters expression of immune related genes. We conclude that Podoplanin expression impacts the immunoregulatory properties of fibroblastic stroma in lymph nodes and is a key transcriptional regulator of fibroblast function in lymph nodes.

*Summary: Podoplanin expression regulates the immunoregulatory properties of fibroblastic stroma in lymph nodes*

## INTRODUCTION

Lymph nodes are key organs for immunity, providing the ideal meeting points for immune cells and coordinating the response to tissue damage and infection(*1*–*3*). Lymph nodes begin to develop at E12.5-E13.5 and are initially composed of both haematopoietic lymphoid tissue inducer (LTi) and stromal lymphoid tissue organiser cells (LTo) cells (*2*, *4*, *5*). The lymph node tissue supports the recruitment of immune cells but can also support their expansion during inflammation(*1*, *6*, *7*). Importantly, in order to accommodate rapidly rising cell numbers, lymph nodes need to reversibly expand (*6*, *8*). Stromal cells constitute a small fraction (1-2%) of the total cells in the lymph node however they are critical for supporting immune cells both during the steady state and disease. Much is now known about the construction of lymphoid tissues through development, and the signals required for commitment and differentiation of specialised lymphoid tissue stroma (*9*–*11*). However, we do not know how mature, adult lymphoid tissues maintain immunoregulatory properties through steady state and inflammatory contexts.

Fibroblastic reticular cells (FRCs) are the most abundant stromal cells of the lymph node forming an interconnected network which supports both the structural integrity of lymph nodes (*12*) and the various immune niches. Recent advances in scRNA-seq have shown high levels of heterogeneity of FRCs and their ability to specifically support distinct niches of immune cells (*9*, *13*). FRCs express the chemokines CCL19 and CCL21 to facilitate the meeting of antigen-presenting dendritic cells and naïve T cells (*13*–*15*). FRCs proximal to high endothelial vessels express high levels ICAM-1 and VCAM-1, important adhesion molecules, supporting the recruitment of both T and B lymphocytes from the circulation (*16*). FRCs proximal to B cell follicles are characterised by their high expression of CCL21 and BAFF to support B cell survival (*17*–*19*). Fibroblastic reticular cells of lymph nodes are the archetype ‘immunofibroblast’ with the power to coordinate adaptive immune responses. As lymph nodes form in embryonic development, mesenchymal precursor cells expand and acquire specialized immunological features to attract and retain immune cells. It is not understood how such a ubiquitous cell type such as a fibroblast adapts to support immune functions.

A common feature of all FRC subsets in the lymph node is their expression of Podoplanin. Podoplanin is a small transmembrane glycoprotein whose functions have been described in different contexts in both health and disease and is important in platelet aggregation, the separation of the lymphatic and endothelial vasculature, and in cell maturation and migration during development (*20*, *21*). Podoplanin is expressed on several cell types including platelets, kidney podocytes (*22*), osteocytes (*23*), alveolar epithelial cells (*24*), keratinocytes (*25*), lymphatic endothelial cells (LECs) (*2*, *26*, *27*) and constitutively expressed by FRCs (*28*). Moreover, Podoplanin levels are upregulated in inflamed tissues (*29*), rheumatoid arthritis (*30*), in cancer cells cancer-associated fibroblasts (*31*, *32*) and during infection (*33*). Podoplanin was first described as a ligand for dendritic cell (DC) migration through the interaction with C-type lectin CLEC-2 (*6*). Later it was shown that this same interaction promotes lymph node expansion through the regulation of FRC contractility and regulates the unilateral deposition of extracellular matrix into the conduit network (*5*–*7*, *34*). Podoplanin maintains actomyosin contractility via activation of the cytoskeletal regulators ezrin and moesin (*35*), two members of the ERM (ezrin, radixin and moesin) family, which tether the cortical actin cytoskeleton to integral proteins in the plasma membrane (*36*). Most recently, we have described Podoplanin as a mechanical sensor in FRCs required to trigger FRC proliferation in response to increasing tissue tension through lymph node expansion (*8*). Mice with full knockout of Podoplanin fail to develop lymph nodes (*37*) and die shortly after birth due to circulatory defects (*38*, *39*), therefore the function of Podoplanin has not been fully explored specifically in FRCs, or in adult lymphoid tissues. Our recent work suggests a more extensive role for Podoplanin in lymph node tissue function, beyond control of actomyosin contractility. We therefore undertook a detailed analysis of how the PDPN/CLEC-2 signalling axis determines transcriptional states in FRCs and directly test the role of Podoplanin in determining the immunoregulatory phenotypes of FRC subsets.

We utilised a mouse model (PDGFRα-mGFP-CreERT2) for conditional genetic manipulation of fibroblasts. Platelet derived growth factor alpha (PDGFRα) was chosen for its broad expression in mesenchymal cells and their progenitors. Further, PDGFRα has been identified as a universal fibroblast marker through extensive single cell transcriptomic analysis of fibroblasts across a range on tissues and inflammatory states (*9*). This model allows us to visualise the fibroblastic reticular network structure and to conditionally delete Podoplanin. In this way we can now assess the function of Podoplanin specifically in fibroblastic stroma of adult lymphoid tissues.

## RESULTS

### Podoplanin deletion *in vivo* alters the transcriptional state of FRCs

We generated a PDGFRα^mGFPΔPDPN^ mouse model to examine the function of Podoplanin on lymph node fibroblasts and lymph node tissue function. This model targets fibroblastic stroma specifically, PDPN^+^ lymphatic endothelial cells (LECs) are not labelled with GFP (Fig. 1A). Tamoxifen treatment of PDGFRα^mGFPΔPDPN^ (ΔPDPN) reduced expression of Podoplanin by >50% in >48.9% of GFP^+^ cells (lymph node fibroblasts) (Fig. 1B).

**Fig. 1.**
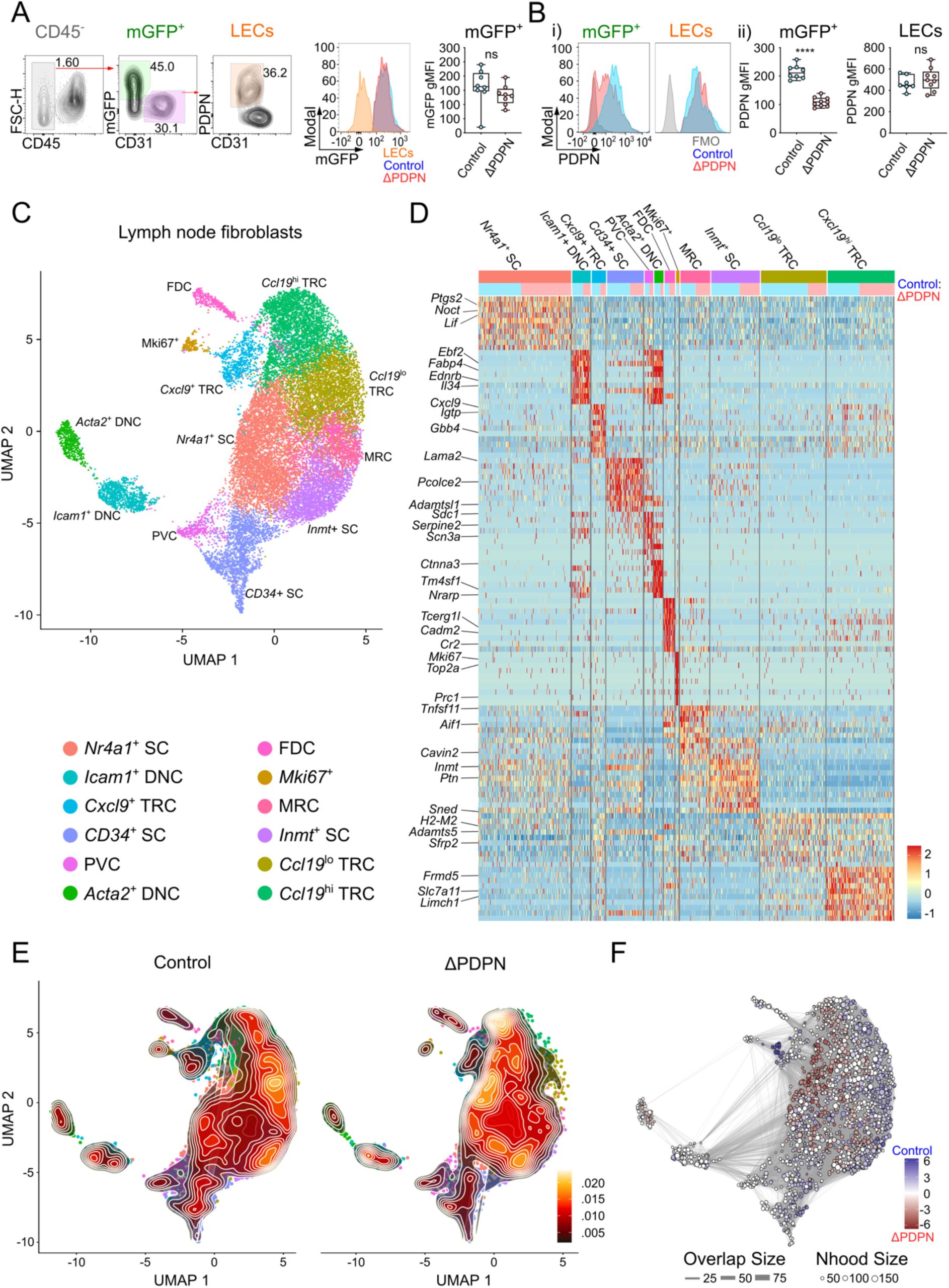
Transcriptional state of fibroblasts is altered by podoplanin deletion. (**A**) Flow cytometry gating of CD45^-^, mGFP^+^ (PDGFRα^mGFP^) and lymphatic endothelial cells (LECs) with representative histograms of the mGFP expression for PDGFRα^mGFP^ (control) or PDGFRα^mGFPΔPDPN^ (ΔPDPN) stroma with geometric mean fluorescence intensity (gMFI) for mGFP^+^. (**B**) i) Representative histograms of PDPN gMFI for mGFP^+^ and LEC populations (ii) gMFI for PDPN in mGFP^+^ (fibroblasts) and LECs. N>8 mice for each genetic model. Mann-Whitney test (two tailed), ****p<0.0001, ns = not significant. (**C**) UMAP plot of fibroblasts (CD45^-^CD31^-^) and (**D**) heatmap showing DEGs. Ratio of Control (blue) and ΔPDPN (pink) numbers (top row). (**E**) Contour density plots of Control versus ΔPDPN fibroblasts (CD45^-^ CD31^-^). (**F**) Neighbourhood distance analysis (MiloR) showing Control (blue gradient) or ΔPDPN (red gradient) enriched areas.

CD45^-^CD31^-^ cells (fibroblasts) were enriched from PDGFRa^mGFP^ (Control) and PDGFRα^mGFPΔPDPN^ (ΔPDPN) lymph nodes 10 days post tamoxifen induction of Cre recombinase (fig. S1A). Cluster analysis identified 12 populations of cells which mapped closely to the subpopulations identified by Rodda et al. (*13*) (Fig. 1,C and D, table S1 to table S3, fig. S1B and fig. S2,A and B). We observed reduced *Pdpn* expression in each of the *Pdgfra^+^* clusters, whereas the *Pdgfrb*^+^ clusters were either *Pdpn* negative or *Pdpn* levels were unaffected by the genetic deletion as expected (fig S2,C and D). In addition, we describe a separate proliferative population of *Mki67*^hi^ FRCs and 2 different populations of double negative cells (DNC) (Fig. 1,C and D, fig. S2B, and tables S1 to S3). The DNC were first described by Malhotra et al. as a contractile pericyte population (*40*), and in concordance with their study we confirm that they express *Acta2* (fig. S1B), *Ednra, Ednrb, Myl9* (fig. S2A), and are a distinct cell type from FRCs since they are *Pdgfrb*^+^ and *Pdgfra*^-^ (Fig. 1,C and D and fig. S1C).

Using these data, we next compared the Control and ΔPDPN stromal populations using density plots (stat_density_2d) (Fig. 1E). To quantify changes to subsets density, we performed neighbourhood analysis using MiloR (*41*) (Fig. 1F). DNC subsets were unaffected which was expected since they do not express *Pdgfra* and therefore are not targeted in our model. However, it is important to note that ΔPDPN FRCs are transcriptionally distinct from these DNC. We observed reduced numbers of FRCs in the *Ccl19*^lo^, *Cxcl9*^+^, and *Cd34*^+^ subsets (blue neighbourhoods) (Fig. 1,D to F). ΔPDPN FRCs clustered more densely in the *Nr4a1* and *Ccl19hi* regions (red neighbourhoods) (Fig. 1,E and F). We performed trajectory analysis (Monocle 3) of the Control and ΔPDPN fibroblasts using progenitor markers CD34 and Pi16 as the initial trajectory point (fig. S2E, white circle). CD34 is a common precursor marker associated with tissue maintenance across multiple organs and Pi16 expression has been described as a marker for mesenchymal precursor cells (*9*, *42*–*44*). The trajectory data shows that the fibroblasts, and especially the *Ccl21^+^*TRC population, lose specificity after deletion of Podoplanin (fig. S2E). Overall, our data show that *Pdpn* expression significantly alters FRC phenotypes across the broad range of subsets as defined by the field.

### ΔPDPN FRCs are transcriptionally distinct from Control FRC subsets

We more closely examined cells clustering in the *Nr4a1* and *Ccl19*^hi^ subsets which were enriched in ΔPDPN lymph nodes. We performed subclustering analysis (Seurat) and found shifts within transcriptional states. For example, within the *Nr4a1* cluster, we found more (80.1:19.9 ratio) ΔPDPN cells in the *Ccl21*^hi^ subcluster and reduced numbers (20.3:79.7 ratio) of ΔPDPN in the *Ccl21*^lo^ subcluster (Fig. 2,A and B, and table S4,).

**Fig. 2.**
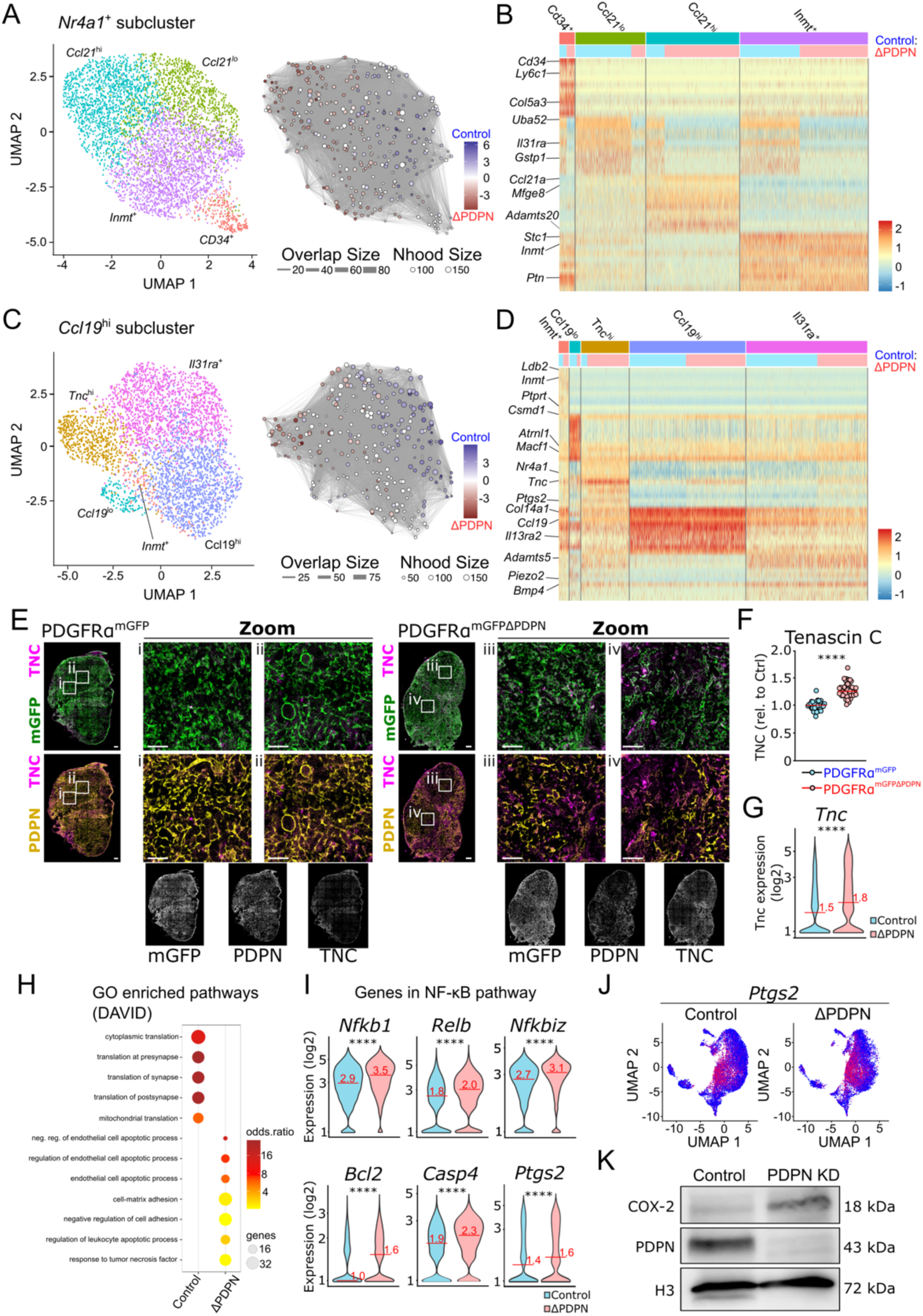
Distinct transcriptional profile in PDGFRα^mGFPΔPDPN^ fibroblasts. (**A**) Subcluster analysis of *Nr4a1*^+^ cluster, miloR showing PDGFRα^mGFP^ (Control -blue gradient) and PDGFRα^mGFPΔPDPN^ (ΔPDPN -red gradient) enriched areas. (**B**) Heatmap showing top 10 DEGs for each defined subcluster, Ratio of Control (blue) and ΔPDPN (pink) numbers (top row). (**C**) Subcluster analysis of *Ccl19*^hi^ cluster and miloR showing PDGFRα^mGFP^ (Control -blue gradient) and ΔPDPN (red gradient) enriched areas. (**D**) Heatmap showing top 10 DEGs for each defined subcluster. Ratio of Control (blue) and ΔPDPN (pink) numbers (top row). (**E**) Tilescans and zoomed areas comparing PDGFRα^mGFP^ and PDGFRα^mGFPΔPDPN^ lymph nodes (mGFP -green), Podoplanin (PDPN – yellow) and Tenascin C (TNC – magenta). (**F**) Quantification of TNC immunofluorescence representative of 4-5 lymph nodes per condition. Scales bar = 50 μm. (**G**) Violin plot showing *Tnc* expression PDGFRα^mGFP^ and PDGFRα^mGFPΔPDPN^ fibroblasts from scRNA-seq analysis. (**H**) GO-enriched pathways (DAVID) in PDGFRα^mGFP^ and PDGFRα^mGFPΔPDPN^ fibroblasts. (**I**) Violin plots with median expression for NF-κB pathway genes: *Nfkb1*, *Relb*, *Nfkbiz*, *Bcl2*, *Casp4* and *Ptgs2*. Mann-Whitney test (two tailed), ****p<0.0001. (**J**) Expression of *Ptgs2* on fibroblast UMAP. (**K**) Western blot for COX-2, PDPN and Histone H3 in FRC cell lines, control versus shRNA KD PDPN (PDPN KD).

When we examined the *Ccl19*^hi^ subset, subclustering analysis revealed an enrichment of ΔPDPN FRCs in a *Tnc*^hi^ subcluster (87.4:12.6 ratio) (Fig. 2,C and D, and table S5). To validate these data, we immunostained lymph nodes from PDGFRα^mGFP^ and PDGFRα^mGFPΔPDPN^ mice and quantified a >30% increase in Tenascin C (TNC) protein expression in PDGFRα^mGFPΔPDPN^ tissues (Fig. 2,E and F) which correlated with the increased overall expression of *Tnc* quantified across all fibroblast subsets in the scRNA-seq data (Fig. 2G). Since Tenascin C is highly linked to inflammatory states (*45*, *46*), we performed DAVID analysis to compare inflammatory pathways between Control and ΔPDPN fibroblasts (Fig. 2H). We found enrichment of biological pathways relating to cell and matrix adhesion and upregulation of many NF-κB target genes in ΔPDPN fibroblasts (Fig. 2,H and I). Interestingly, NF-κB signalling, downstream of lymphotoxin-β receptor engagement, has been implicated in FRC development (*47*). *Ptgs2,* encoding COX-2, a key enzyme to produce inflammatory prostaglandins (*48*), is positively regulated by NF-κB and is enriched in ΔPDPN fibroblasts (Fig. 2,I and J). COX-2 expression by FRCs has been linked suppression of T cell activation in lymph nodes (*49*). We tested whether Podoplanin impacted COX-2 expression at protein level in FRC cell lines *in vitro,* using shRNA to knockdown PDPN expression, and confirmed by western blotting upregulation of COX-2 in PDPN KD cell lines (Fig. 2K). We therefore link Podoplanin directly and intrinsically to NF-κB signalling and COX-2 expression in FRCs.

### Lymph node network integrity and FRC inflammatory profile altered by Podoplanin

Since our analysis highlighted the cell-matrix adhesion GO term as enriched in ΔPDPN FRCs (Fig. 2H), we next asked if other matrix components were altered in PDGFRα^mGFPΔPDPN^ lymph nodes. Tilescans showed increased deposition of perlecan, a basement membrane component, in PDGFRα^mGFPΔPDPN^ lymph nodes (Fig. 3A) which confirmed that matrix remodelling is widely impacted by Podoplanin deletion. We observed areas of the paracortex tightly packed with leukocytes (nuclei stained with DAPI) but devoid of GFP^+^ fibroblasts in PDGFRα^mGFPΔPDPN^ mice. These areas also lacked extracellular matrix components of the conduit network suggesting that the reticular network had lost integrity (Fig. 3A). To quantify the disruption of the FRC network in the paracortex we used an ImageJ tool (Mitochondrial Network Analysis - MiNA) based on ‘skeletisation’ to quantify mathematical properties of the network (Fig. 3,B and C) (*50*). We find in PDGFRα^mGFPΔPDPN^ mice there is a 60% decrease in the number of branches, shortened branch length and a 25% decrease in the network footprint, confirming a severe disruption to the FRC network (Fig. 3,B and C).

**Fig. 3.**
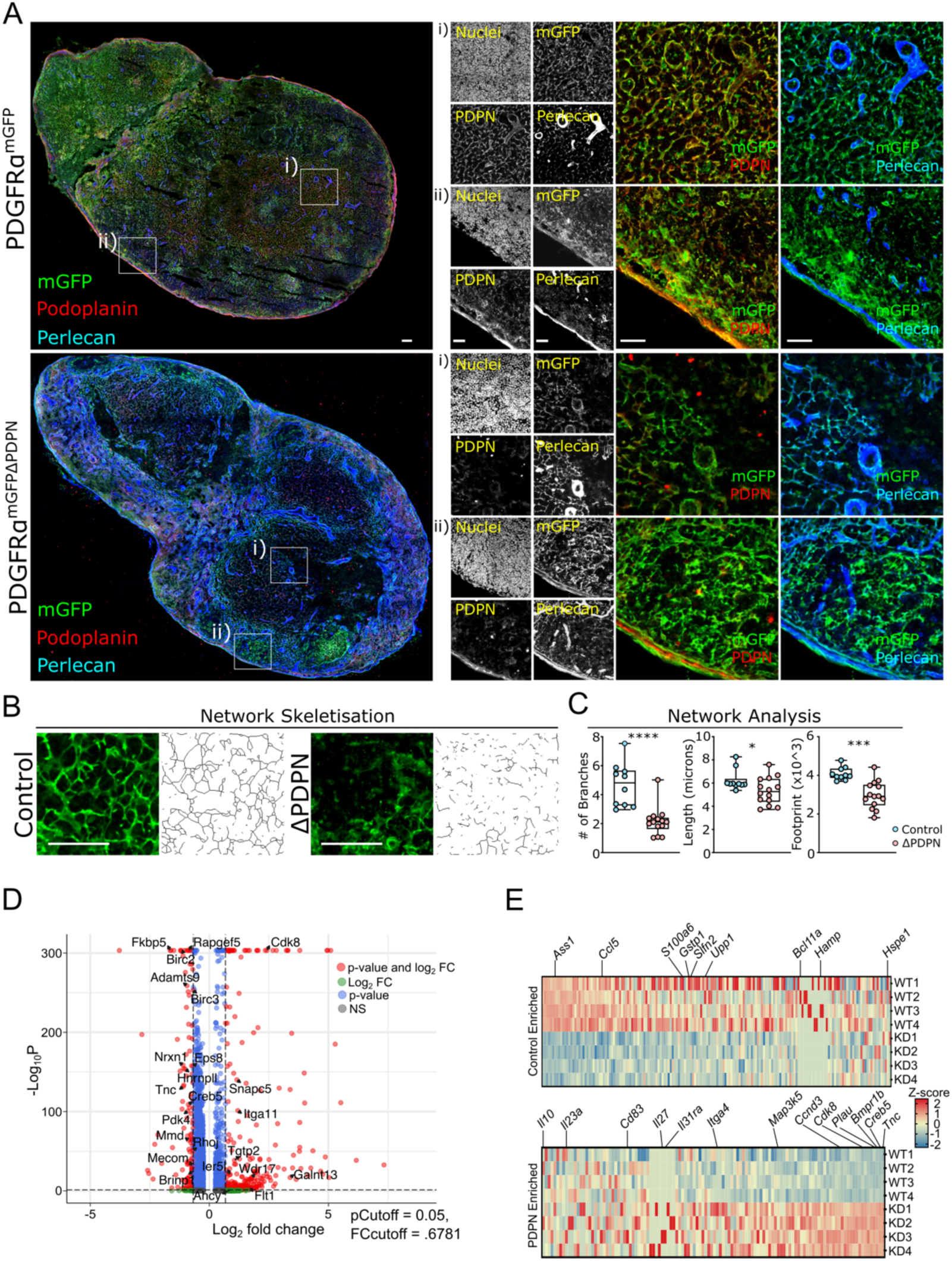
Lymph node network integrity and FRC inflammatory profile altered by Podoplanin. **(A)** Representative tilescan of PDGFRα^mGFP^ (Control) or PDGFRα^mGFPΔPDPN^ (ΔPDPN) lymph nodes. Staining for Podoplanin (red), mGFP (green) and perlecan (blue); zoomed images of (i) paracortex (T cell area) (ii) capsule. Scale bar = 50 μm. **(B)** Representative images of FRC network (mGFP) and network skeletisation for PDGFRα^mGFP^ (control) or PDGFRα^mGFPΔPDPN^ (ΔPDPN) lymph nodes. Scale bar = 100 μm. **(C)** Network analysis quantifying the number of network branches, length of branches and network footprint. Network quantification N=10-15 ROI of lymph node paracortex; Mann-Whitney test (two tailed), ****p<0.0001, ***p<0.001, *p<0.05. **(D)** Volcano plot showing genes significantly altered after PDPN deletion (scRNA-seq dataset). **(E)** DEGs from (D) overlaid with *ex vivo* bulk RNA-seq of FRC cell lines expressing basal levels control versus shRNA KD PDPN (PDPN KD).

The scRNA-seq data provide evidence that Podoplanin expression by FRCs is important for determining their transcriptional state, which impacts their subset identity (Fig. 1 and 2). However, the transcriptional changes *in vivo* could be caused by a combination of cell intrinsic mechanisms and by alterations in crosstalk between other tissue cell types. To address how the Podoplanin intrinsically regulates gene expression in FRCs, we silenced Podoplanin expression in FRC cell lines *in vitro* and performed bulk RNA-seq. This dataset provides a robust, reproducible readout of average cell state therefore highlights genes regulated by Podoplanin which are not dependent on the tissue microenvironment. We compared a list of 403 genes significantly altered by Podoplanin expression across all *in vivo* cell clusters to our bulk RNA-seq analysis (Fig. 3D). Of these, 290 were also found in our the Podoplanin -dependent gene signatures *in vitro* (Fig. 3E, and table S6). Interestingly, many of these genes directly regulated by Podoplanin expression were targets of NF-κB as predicted by the DAVID analysis of our *in vivo* data (Fig. 2H). Of note, *Tnc* was upregulated across multiple stromal subsets in the scRNA-seq data, validated in tissue staining at protein level and indeed also upregulated in PDPN-KD FRC cell lines *in vitro*. Upregulation of *Plau* (Fig. 3E), another gene inversely correlated with Podoplanin expression, has been linked conversion of fibroblasts to inflammatory-type cancer associated fibroblasts in oesophageal SCC (*51*). On the other hand, downregulation of *Ass1* (Fig. 3E) is linked to pulmonary fibrosis (*52*). We therefore conclude that Podoplanin can directly and intrinsically impact the inflammatory state of FRCs.

### PDPN/CLEC-2 signalling induces two distinct transcriptional signatures

While we show that Podoplanin can directly change the phenotype of FRCs *in vivo* and *in vitro*, FRCs also have important crosstalk with neighbouring immune cells. Podoplanin is a membrane glycoprotein that can bind and cluster with CLEC-2 (*53*), a C-type lectin expressed on migratory DCs, platelets and other myeloid cells (*27*, *54*). We exploited the *in vitro* FRC cell lines to directly test the effect of CLEC-2 engagement and compare immediate (6 hour) and longer-term (24 hours) transcriptional changes to FRCs (Fig. 4). Using this experimental set up we were able to isolate genes that were intrinsically Podoplanin-dependent and importantly were unaffected by ligand binding (Fig. 4A, and table S7). The Podoplanin-dependent upregulated genes included fibroblast markers (*Thy1*), vesicular transport and autophagy (*Atg3, Atg12, Pink1, Lamp2, Dnm1, Tapb2*), cell signalling (*Jak2, Gatsl3, Dusp16*), actin dynamics (*Rnd1, Ssh3*), and transcription factors (*Sox11, Ahr, Hoxd9, Cxxc5*) (fig. S3A).

**Fig. 4.**
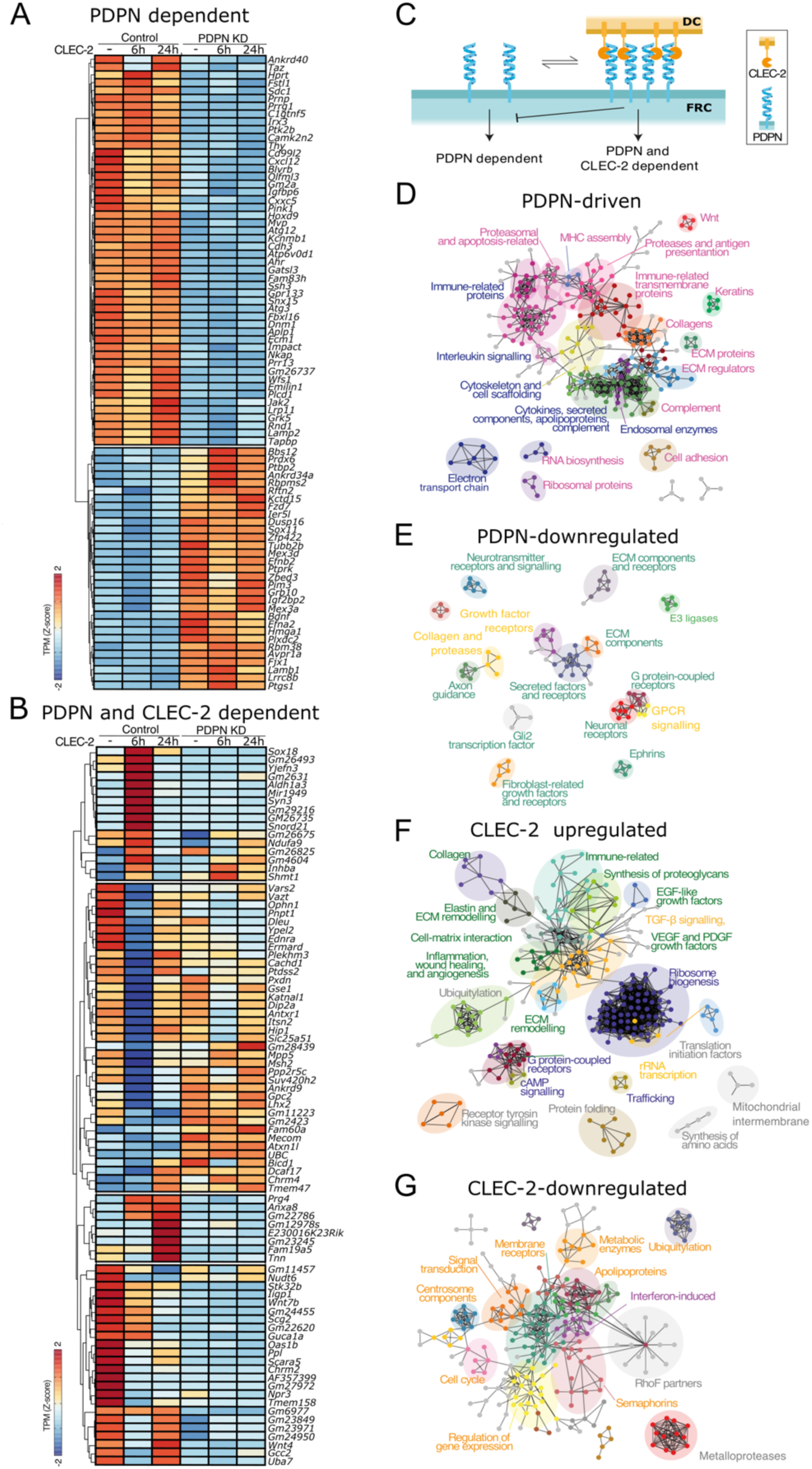
PDPN/CLEC-2 signalling axis controls the transcriptional landscape of FRCs *ex vivo*. (**A**) Heatmap of the genes regulated by PDPN expression independently of CLEC-2 signalling. (**B**) Heatmap of the genes regulated by CLEC-2 in PDPN^+^ (Control) FRC cell line. (**C**) Diagram of proposed modes of transcriptional regulation. (**D**) STRING analysis of PDPN-driven, (**E**) PDPN-downregulated, (**F**) CLEC-2 upregulated and (**G**) CLEC-2 downregulated genes.

When we examined the genes transcriptionally regulated after treatment with CLEC-2 (Fig. 4B, and table S7), we observed most differentially expressed genes occurred at 6 hours (Fig. 4B). This finding supports previous knowledge that CLEC-2 signalling is transient in FRCs (*34*). It has been previously reported that signalling of Podoplanin to cytoskeletal modulators is dampened by binding of CLEC-2 (*6*) and exposing FRCs to CLEC-2 phenocopies a loss of Podoplanin through reduction in actomyosin contractility (*6*). Therefore, we asked whether CLEC-2-upregulated genes correlated with PDPN KD. Hierarchical clustering revealed four PDPN and CLEC-2-dependent groups (Fig. 4B). The two largest clusters were downregulated by CLEC-2 at 6 hours (including the Wnt ligands *Wnt4* and *Wnt7b*, and the extracellular matrix components *Tnn* and *Prg4*) and only a small fraction of these genes followed the previously reported pattern phenocopying PDPN KD (Fig. 4B). We found only a small fraction of hits (85 and 236 genes), overlap between the genes regulated by Podoplanin or CLEC-2 (fig. S3A) suggesting that while in some cases CLEC-2 binding does phenocopy loss of Podoplanin expression, CLEC-2 also induces a distinct transcriptional signature (Fig. 4C).

We used the STRING functional annotation platform to study functional networks, including physical interactions as well as functional connections (Fig. 4,D to G). PDPN-driven genes formed a large cluster of immune-related genes (Fig. 4,D and E, and table S8). This cluster included membrane receptors (*Cd47*, *Itgb2*), cytokine signalling (*Stat1*, *Stat2*, *Irf7*, *Fit1*, *Ifit1*, *Ifit3*, *Cxcl1, Cxcl10, Cxcl5, Ccl2, Ccl9*), several collagen genes, ECM components (*Ecm1, Timp3*) consistent with early microarray analysis of lymph node stromal cells (*40*), complement genes (*C2, C3, C1ra, C1s1, C1s2*), and genes related to antigen presentation (*Anpep, H2-T23, H2-K1*). CLEC-2 regulated genes formed a large cluster of hits related to ribosome biogenesis and regulation of translation (Fig. 4,F and G, and table S8). Moreover, in line with the PDPN-driven genes, we found clusters associated to immune-related functions and signalling, including different families of growth factors (*Tgfb*, *Vegf*, *Pdgf*), genes involved in inflammation, wound healing, and angiogenesis. Moreover, PDPN expression and CLEC-2 binding altered expression of many transcription factors which would be expected to have wide ranging effects on the transcriptional profile of FRCs (fig. S3,B and C). We performed gene-enrichment analysis using the GO Biological Process (http://genontology.org) database and found that PDPN-driven pathways included response to mechanical stimuli and extracellular matrix organisation as have been previously reported (*8*, *34*) (fig. S3,D to G), but which we can now also confirm *in vivo* in our scRNA-seq analysis (Fig. 2).

### Stromal/immune cell crosstalk is altered in PDGFRα^mGFPΔPDPN^ lymph nodes

To examine the impact of stromal PDPN deletion on immune cell crosstalk in tissues we quantified immune cell populations in PDGFRα^mGFP^ and PDGFRα^mGFPΔPDPN^ lymph nodes (fig. S1B). CD45^+^ and CD31^+^ cells (immune and endothelial cells) were enriched from Control and PDGFRα^mGFPΔPDPN^ lymph nodes (fig. S1B). We identified 21 clusters of immune and endothelial cells with distinct transcriptional profiles (Fig. 5A and B, fig. S4, table S9). Neighbourhood analysis by miloR showed a loss of myeloid cells, macrophages, B cells and CD4^+^ T cells in PDGFRα^mGFPΔPDPN^ (blue neighbourhoods) (Fig. 5C).

**Fig. 5.**
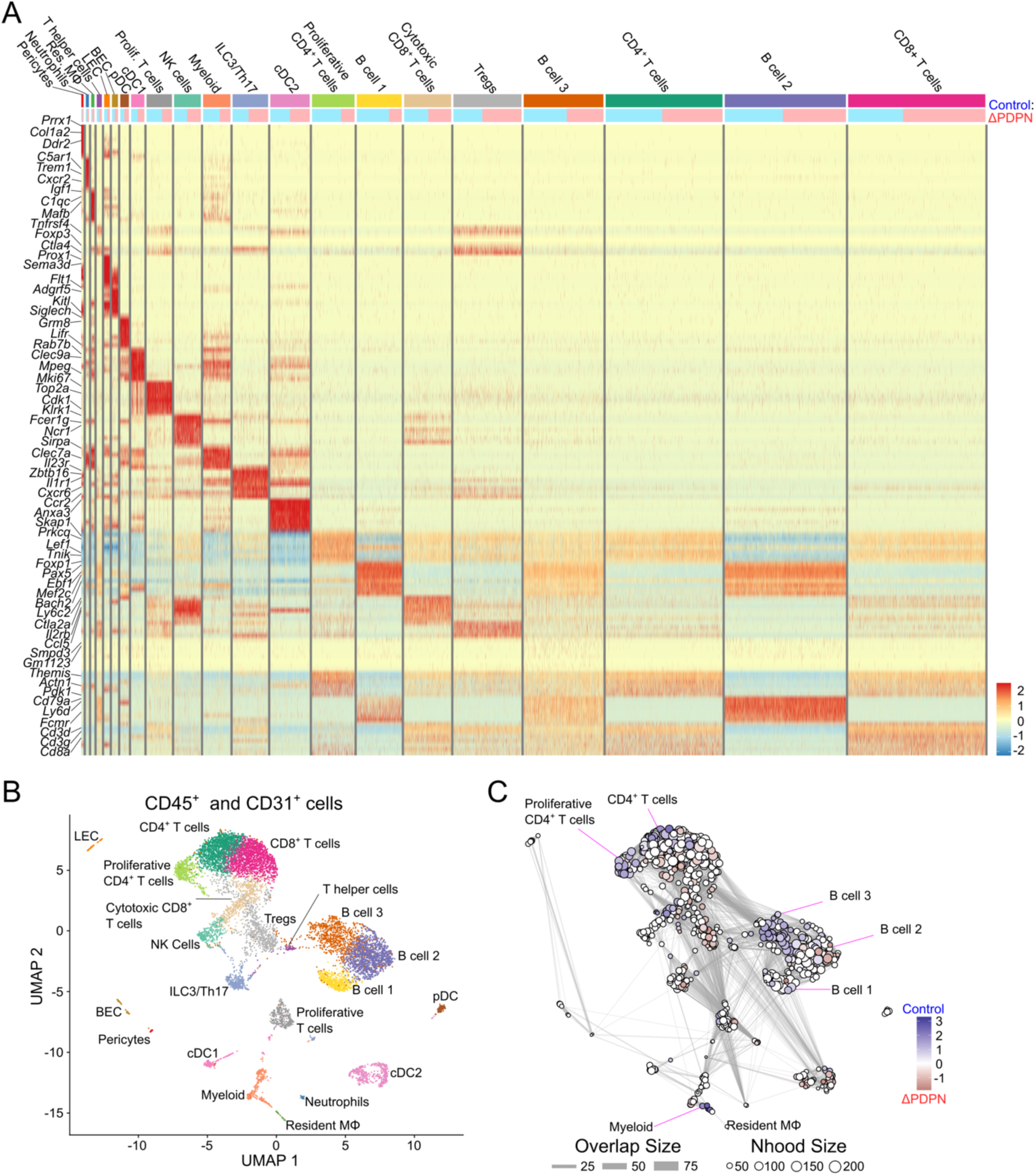
Immune cell transcriptional profile is altered by Podoplanin deletion. (**A**) Heatmap analysis of immune and endothelial clusters (**B**) UMAP and (**C**) MiloR neighbourhood analysis showing Control (blue gradient) and ΔPDPN (red gradient) abundant areas.

### Stromal cross talk with innate immune cells is reduced in the absence of Podoplanin

To understand how the crosstalk between immune cells and stromal cells is affected we performed CellChat analysis between stromal and immune populations (fig. S5) (*55*, *56*). Many of the enriched pathways in the PDGFRα^mGFPΔPDPN^ lymph nodes support a proinflammatory profile, including RANKL, TENASCIN, TGFb, and CXCL (fig. S5A). We examined both the number of interactions (fig. S6B), and the interaction strength (fig. S5C) between stromal cells and immune cells. Our data show that interactions between stromal cells and myeloid cells were particularly attenuated in PDGFRα^mGFPΔPDPN^ lymph nodes; whereas interactions between stromal cells and cDC2 cells were actually enhanced after Podoplanin deletion (fig S5B and C).

Past studies have also shown the importance of CLEC2:PDPN signalling in driving dendritic cell migration and lymph node activation during immune responses (*6*). We also find CLEC-2 dependent transcriptional regulation through PDPN (Fig. 4). We therefore performed subcluster analysis on the myeloid and resident macrophage populations, and dendritic cells (Fig. 6,A and B, and tables S11 and 12). We observed a reduction in the number of inflammatory monocytes/macrophages in PDGFRα^mGFPΔPDPN^ lymph nodes, however there was an enrichment in the monocyte derived macrophage cluster (red gradient) (Fig.6,A and B). We also performed subcluster analysis on all DC clusters: cDC1, cDC2 and pDCs (Fig. 6,C and D, and table S12). Neighbourhood analysis revealed shifts in within the cDC2 subclusters with *Ccl5*^hi^ cDC2 being enriched in Control tissues (Blue) whereas *Nrp2*^+^ cDC2 were enriched in PDGFRα^mGFPΔPDPN^ tissues (Red) (*57*). To validate these transcriptomic data, we examined myeloid cells by flow cytometry and microscopy (Fig. 6,E and F, and fig. S6). In PDGFRα^mGFPΔPDPN^ lymph nodes we observed a reduction of CD103^+^ DC and but an increase in the percentage of Ly6c^hi^ Monocytes (Fig. 6D). We compared numbers and location of CD11b^+^ cells in lymph node tissue sections and observed reduced CD11b^+^ cells in PDGFRα^mGFPΔPDPN^ lymph nodes (Fig. 6,G and H). The CD11b^+^ cells in PDGFRα^mGFPΔPDPN^ lymph nodes were also more circular, which is consistent with the increased number of monocytes (Fig. 6,G and H). Overall, this suggests that in addition to affecting the transcriptional state of fibroblasts, PDPN deletion also affects the activation state and cellular composition of myeloid cells in the lymph node at steady-state.

**Fig. 6.**
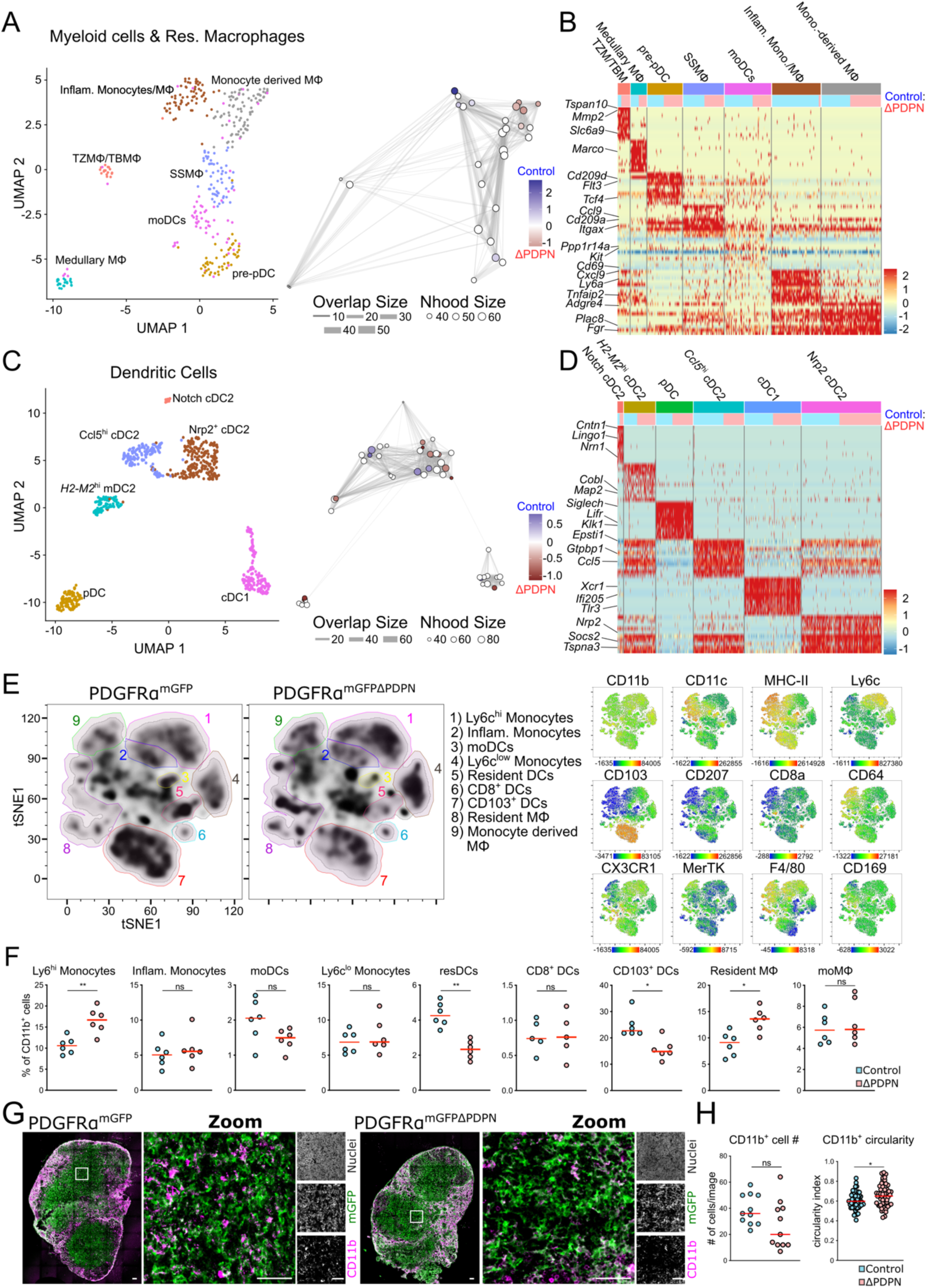
Innate cell phenotype and numbers altered in PDGFRα^mGFPΔPDPN^ lymph nodes. (**A**) Subcluster analysis of Myeloid & Resident Macrophages showing UMAP, MiloR neighbourhood distance analysis (**B**) Heatmap of DEGs of Myeloid & Resident Macrophages. **(C)** Subcluster analysis of Dendritic cells (cDC1, cDC2 and pDC clusters) showing UMAP, MiloR neighbourhood distance analysis **(D)** Heatmap DEGs for dendritic cell populations. **(E)** Flow cytometry analysis of innate immune cells (CD45^+^CD11b^+^), tSNE plots using the markers: CD11c, I-A/I-E (MHC-II), Ly6c, CD103, CD207, CD8a, CD64, CX3CR1, MerTK, F4/80 and CD169. (**F**) Changes in the percentages of each innate cell population comparing Control and ΔPDPN. Mann-Whitney test (two tailed), ns = no significance, **p<0.01, *p<0.05. (**G**) Tilescans of PDGFRα^mGFP^ and PDGFRα^mGFPΔPDPN^ lymph nodes stained for mGFP (green) and CD11b (magenta) **(H)** Quantification of number of CD11b^+^ cells per ROI and circularity index of each CD11b^+^ cell. Scale bar = 50 μm. Images representative of 4-5 lymph nodes for each genetic model Mann-Whitney test (two tailed), ns = no significance, *p<0.05

### Disrupted stromal-immune crosstalk in PDGFRα^mGFPΔPDPN^ mice attenuates B cell activation

In addition to reduced crosstalk with innate cells, we also observed changes to B cell populations in PDGFRα^mGFPΔPDPN^ lymph nodes (Fig. 5). We performed subcluster analysis of B cell populations to understand how Podoplanin deletion affects their phenotype (Fig. 7A, and table S12). MiloR analysis shows altered neighbourhood clusters and there were fewer activated or germinal centre-like B cells in PDGFRα^mGFPΔPDPN^ lymph nodes (Fig. 7,A and B). B cell follicles are supported by a stromal network of FDCs which are a distinct subset of stromal cells that specifically support B cells through the production of CXCL13 for their recruitment, and growth factors such as BAFF (*13*, *17*). FDCs also form the stromal scaffolds for germinal centre (GC) formation and retain and present antigen to B cells to support B cell selection (*58*). Since FDCs are of mesenchymal origin and express PDGFRα (*13*) they also conditionally targeted in our PDGFRα^mGFPΔPDPN^ mice. FDCs also express PDPN (fig. S2), although at lower levels than FRCs, but the function of PDPN in FDCs has not previously been investigated. To understand why B cell populations would be affected by Podoplanin deletion, we next performed subcluster analysis of the stromal cells supporting B cell function, FDCs (Fig. 7C) and MRCs (Fig 7D), (fig. S7,A and B, and tables S13 and S14). In PDGFRα^mGFPΔPDPN^ lymph nodes miloR analysis highlighted shifts in FDCs cluster towards reduced expression of *Fcgr2b*, which would be consistent with reduced activation of B cells (Fig. 7A, fig. S7A, and table S13). In tissues, we measured the area covered by the FDC network in each B cell follicle (fig. S7C) and found a similar number of B cell follicles and FDC network size between PDGFRα^mGFP^ and PDGFRα^mGFPΔPDPN^ mice. However, the follicle size was slightly reduced in the latter (fig. S7C). The MRC subcluster analysis also revealed a shift towards MRCs expressing *Nr4a1* in PDGFRα^mGFPΔPDPN^ mice (Fig. 7D, fig. S7B, and table S14). Interestingly, the PDGFRα^mGFPΔPDPN^ MRC populations also showed higher expression of inflammatory pathways such as *Ptgs2, Cxcl1,* similarly to TRC populations (fig. S7D, and Fig. 2). Overall, these findings show that Podoplanin drives changes in the phenotype of FDCs and MRCs which could compromise the crosstalk with B cells.

**Fig. 7.**
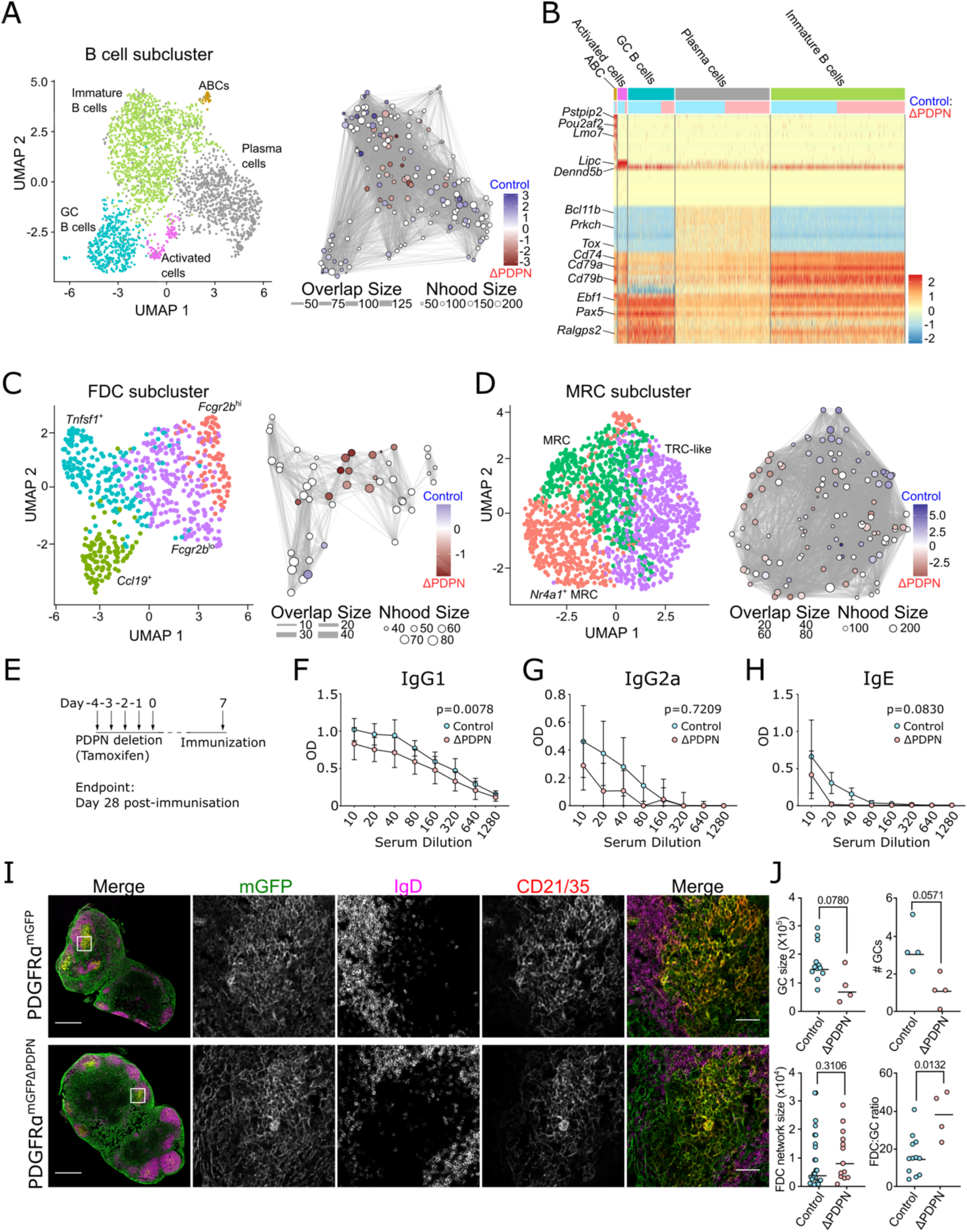
B cell function during steady-state and inflammation compromised by Podoplanin deletion. (**A**) B cell subcluster UMAP and MiloR showing PDGFRα^mGFP^ (Control -blue gradient) and PDGFRα^mGFPΔPDPN^ (ΔPDPN -red gradient) enriched areas. (**B**) Heatmap showing top 10 DEGs for each defined subcluster, ratio of Control (blue) and ΔPDPN (pink) numbers (top row). (**C**) FDC and (**D**) MRC subcluster UMAP and MiloR showing PDGFRα^mGFP^ (Control -blue gradient) and ΔPDPN (red gradient) enriched areas. (**B**) Heatmap showing top 10 DEGs for each defined subcluster, ratio of Control (blue) and ΔPDPN (pink) numbers (top row). (**E**) Schematic of experimental design for immunisation (IFA-OVA) after Podoplanin deletion. Ovalbumin-specific (**F**) IgG1, (**G**) IgG2a and (**H**) IgE in serum from PDGFRα^mGFP^ (Control) and PDGFRα^mGFPΔPDPN^ (ΔPDPN). Wilcoxon test, p-values indicated. (**I**) Tilescans of PDGFRα^mGFP^ and PDGFRα^mGFPΔPDPN^ lymph nodes stained for mGFP (green), IgD (magenta) and CD21/35 (red). (**J**) Analysis of germinal centre (GC) size, GC number FDC network size and ratio of FDCs to GCs. Mann-Whitney test (two tailed), p-values indicated. Scale bar = 500 μm and 50 μm in zoomed images. Images representative of 4-5 lymph nodes for each condition from 2 independent experiments.

The defect in B cell activation could be a direct intrinsic defect, a failure of T cell help or insufficient stromal cell support. We therefore examined if the deletion of PDPN alters adaptive immune responses *in vivo* after immunisation. We compared early activation (day 5) and later stages (day 9) (fig. S8). PDGFRα^mGFPΔPDPN^ lymph nodes expanded less than controls (fig. S8). We have shown previously that PDPN is a mechanical sensor in FRCs and is required to trigger FRC proliferation in response to increased mechanical strain as the lymph node begins to expand (*8*). To specifically quantify how B cell activation is affected by Podoplanin deletion we also measured antibody responses in the serum 28 days post immunisation. We found that PDGFRα^mGFPΔPDPN^ mice had reduced levels of antigen (ovalbumin) specific IgG1, IgG2a and IgE in the serum (Fig. 7,E to H), thereby confirming that Podoplanin deletion has a significant functional impact on adaptive immune responses. Tissue analysis of immunised mice showed there is a trend towards fewer GCs per lymph node in PDGFRα^mGFPΔPDPN^ mice, and those GCs present tend to be smaller (Fig. 7,I and J). This results in altered structure of GCs in PDGFRα^mGFPΔPDPN^ mice as measured by an increased ratio (>2 fold) of FDC to GC area (Fig. 7J) suggesting that there may be a difference in the functional capacity of FDC in PDGFRα^mGFPΔPDPN^ LNs to support B cell activation and that PDPN also plays a role in FDC specification and function.

Overall, our data show that Podoplanin regulates the transcription profile of FRCs in addition to controlling cellular mechanics and actomyosin contractility. Loss of Podoplanin broadly upregulates expression of inflammatory molecules and pathways which will impact many different immune cell types and the overall immunoregulatory functions of the lymphoid tissues.

## DISCUSSION

Podoplanin has complex functions which until now have only been partially described. Podoplanin was first described as a ligand for DC trafficking (*27*) but has since been shown to also maintain HEV integrity through crosstalk with platelets (*54*), regulate extracellular matrix deposition (*34*) and to control the contractility (*6*, *7*) and tissue tension (*8*) of the FRC network. Podoplanin acts as a mechanical sensor in FRCs to trigger proliferation and lack of Podoplanin in FRCs attenuates the acute phase of lymph node expansion (*8*). Now, we show that Podoplanin regulates FRC transcriptional programmes and cell state in both tissue contexts and intrinsically in simple *in vitro* cell systems. Broadly, loss of Podoplanin expression induced upregulation of the NF-κB signalling pathway and expression of pro-inflammatory molecules. For example, COX-2 driving prostaglandin expression (*59*, *60*) and Tenascin C which is a proinflammatory matrix protein normally associated with wound healing, myofibroblast activation and fibrosis (*45*, *46*). In the lymph node, these transcriptional shifts led to reduced numbers of *Cxcl9*^+^, *Ccl19*^+^ and *Ccl21*^+^ TRC subsets and an enrichment of FRCs expressing *Nr4a1*, a subset previously associated with early activation markers (*13*). This pro-inflammatory change in inflammatory state *in vivo* may have been caused by crosstalk between stromal cells and immune cells, however a significant proportion of this transcriptional shift was also observed in bulk RNA-seq analysis *in vitro* using immortalised FRC cell lines, meaning that Podoplanin was directly and intrinsically affecting the phenotype of lymph node fibroblasts. We further tested how the PDPN/CLEC-2 signaling axis impacted the immunoregulatory properties of FRCs *in vitro* and *in vivo,* which revealed that Podoplanin expression in fibroblastic stroma instructs a second, ligand-dependent, transcriptional programme that drives extensive alterations in gene expression in FRCs.

Podoplanin-dependent FRC phenotypes impacted immune cell function in steady state and in reactive lymph nodes. In PDGFRα^mGFPΔPDPN^ lymph nodes the crosstalk between stromal cells and myeloid cells such as macrophages and monocytes were negatively impacted. We found reduced numbers of myeloid cells and dendritic cell subsets in PDGFRα^mGFPΔPDPN^ lymph nodes by flow cytometry and tissue staining. Interestingly, B cell function was also dependent on Podoplanin expression on lymph node stroma (FDC and MRC). We found structural changes to B cell follicles in PDGFRα^mGFPΔPDPN^ lymph nodes in steady state which then impacted B cell activation, germinal centre formation and antibody class switching. These data suggest that Podoplanin has important roles in FDCs in additional to TRCs which has not previously been reported.

Podoplanin is expressed by lymphoid tissue fibroblasts from early in development. The interaction between CLEC-2 and Podoplanin is required for the development and maintenance of lymph nodes (*39*). Indeed, lymph nodes do not develop in a full knockout of Podoplanin (*37*). In contrast, FRCs in aged lymph nodes exhibit impaired upregulation of Podoplanin making them less able to respond to immune challenges (*61*). Beyond secondary lymphoid tissues, PDPN positive stromal cells are key for the early support of tertiary lymphoid structure development (*62*) building the case that Podoplanin is essential for the immunoregulatory properties of lymphoid tissue associated stroma at all life stages.

Podoplanin expression in lymph node stroma is notably altered in a variety of pathological states. For example, lymph nodes from diffuse large B cell lymphoma (DLBCL) patients show upregulation of Podoplanin which is associated with increase stromal density, immunosuppression of CD8^+^ cytotoxic T cells and immune evasion of lymphoma B cells (*63*). This change in stromal phenotype correlated with reduced adaptive immune function, and a decrease in germinal centre formation.

Fibroblastic stroma is also ubiquitous across diverse tissues (*9*). Fibroblast populations are specifically adapted to the functional requirements of their tissue microenvironments (*42*, *64*, *65*). Fibroblasts are also remarkably plastic, able to change their phenotype and function in response to inflammation or tissue damage (*65*, *66*). Here we have studied Fibroblastic reticular cells of lymph nodes which are the archetype immunoregulatory fibroblast. However, how fibroblasts such as TRCs acquire immunoregulatory functions is not well understood (*2*, *3*). Immunoregulatory fibroblasts can also exist outside of lymphoid tissues. For example, both mouse and human tissue studies indicate that some cancer-associated fibroblasts acquire inflammatory phenotypes and can alter immune function in tumours (*66*). PDPN^+^ fibroblasts also spontaneously develop in a wide variety of inflammatory conditions (*21*). Our data confirms the importance of Podoplanin in the functional phenotype of FRCs, and the role of Podoplanin in inflammatory transcriptional signatures. These data lead us to propose that Podoplanin expression is fundamental to FRC function in lymph nodes and may determine the immunoregulatory properties of fibroblasts more widely across other tissues and pathologies.

## MATERIALS AND METHODS

### Mice

Experiments were performed in accordance with national and institutional guidelines for animal care and approved for the Laboratory for Molecular and Cell Biology by the institutional animal ethics committee review board, European research council, and the UK Home office. Breeding of animal lines were maintained at Charles River Laboratory. For generation of the *Pdgfra*-mGFP-CreERT2 and *Pdgfra*-mGFP-CreERT2 x *Pdpn*^fl/fl^ refer to Horsnell et al., 2022 (*8*). Females and males aged between 8 and 13 weeks were used for experiments, unless stated otherwise.

### Tamoxifen administration

Activation of the Cre recombinase was achieved through the administration of tamoxifen (22 mg ml−1; Sigma-Aldrich) resuspended in corn oil (Sigma-Aldrich) or sunflower oil (Sigma-Aldrich). Tamoxifen was dosed (82 μg g−1) intraperitoneally on 3 consecutive days or by gavage for 5 consecutive days. For immunisation experiments, mice were immunised 7 days after tamoxifen treatment. For steady-state experiments mice were sacrificed 10 days after tamoxifen treatment. Inguinal and axillary lymph nodes were studied via flow cytometry or confocal microscopy. Steady state analysis is shown as 4-6 mice, representative of at least 3 independent experiments.

### Immunizations

*In vivo* immunisations were performed with an emulsion Incomplete Freund’s Adjuvant (IFA) containing SIINFENKL (OVA) (Hooke Laboratories). Mice were injected with 100 μl of the emulsion subcutaneously, 100 μg OVA per mouse, on the right flank at the height of the hip 7 days after receiving the final administration of tamoxifen. Animals were sacrificed at 5 and 9 days after immunisations and paired analysis of the inguinal and axillary lymph nodes was performed via flow cytometry and confocal microscopy respectively. For B cell responses animals were sacrificed 28 days post-immunisation and serum and lymph nodes were collected. All *in vivo* data immunisation experiments 3-5 mice were used per experiment and data is pooled or representative of 3 independent experiments.

### Flow cytometry

Lymph node single cell suspension of 2.5×10^6^ cells was incubated for 20 min at 4°C with a purified rat IgG2b anti-mouse CD16/CD32 receptor antibody (BD Biosciences). For stromal cells and lymphocyte staining cells were stained fluorochrome-conjugated antibodies against CD45, CD31, PDPN, CD8a, CD3, CD19, CD4; for innate cell analysis cells were stained for CD11b, B220, CD11c, Ly6C, Ly6G, MerTK, CX3CR1, F4/80, CD169, I-A/I-E, CD207, CD8a, CD64 and CD103 for 25 min at 4°C in PBS containing 1% BSA, 5 mM EDTA, 0.05% NaN3. Cells were incubated with fixable viability stain for 30 min at 4°C. Samples were analysed on BD LSRFortessa x-50 (FACS Symphony) equipped with 100-mW 405-nm, 100-mW 488-nm, 150-mW 561-nm, 100-mW 637-nm and 60-mW 355-nm lasers with a ND1.0 filter in front of the FSC photodiode. Data was collected on the FACSDiva software (version 7).

### Flow cytometry analysis

Flow cytometry data was analysed using FlowJo Software (Tree Star). For tSNE analysis of innate cells, samples were prepared by concatenating approximately 18,000 cells of CD11b^+^ cells per experimental model. tSNE analysis was performed using the FlowJo package and selected parameters: CD11c, CD103, Ly6c, CD64, CD103, CD8a, F4/80, CD169, MerTK, CX3CR1, MHC class II and CD207. The tSNE package settings were set as: Iterations 1000, perplexity 30, learning rate: 1266; KNN algorithm: Exact; gradient algorithm: Barnes-Hut. Cell groups were defined by the expression of the functional markers used for tSNE analysis.

### Immunofluorescence of tissue sections

Lymph nodes placed in Antigen fix (DiaPath) and incubated for 2 hours on ice with gentle agitation. Lymph nodes were washed with ice cold PBS and placed in 30% w/v sucrose containing 0.5% NaN3 and incubated at 4°C overnight. The LNs were placed into Tissue-Tek optimum cutting temperature compound and then embedded into moulds containing the optimum cutting temperature compound. Samples were sectioned on the Leica cryostat (CM1950) at a thickness of 15 μm.

For immunofluorescence tissue sections were permeabilised and blocked for 2 hours at room temperature with 10% normal goat serum (NGS), 0.3% Triton X-100in PBS. Primary antibodies were prepared in the dilutions described in Supplementary Table 3 in PBS containing 10% normal goat serum and 0.01% Triton X-1000 and then centrifuged at 15,000g for 5 minutes. The antibody cocktail was added to the tissue sections which were incubated overnight at 4°C. Sections were then incubated at room temperature for 1 hour before being washed with PBS-0.05% Tween 20 (3 x 15 minutes at room temperature). Secondary antibodies were prepared similarly to the primary antibodies and added to the tissue for 2 hours at room temperature. Sections were then washed with PBS-0.05% Tween 20 (2 x 15 minutes), PBS (1 x 10 minutes) and H_2_O (1 x 5 minutes) at room temperature and were then mounted using mowiol mounting media. Images of the tissue were obtained on the Leica TCS SP5 with HCX Plan-Apochromat x40 (NA 1.25) oil lenses. Images were captured at 1,024×1,024 pixels, three-line and 2-frame average onto hybrid pixel or photomultiplier tube detectors. Imaging areas were manually detected, and z-stacks (15-25 μm) were acquired with intervals at 0.3-0.6 μm. Tile scan boundaries were manually set, and images were stitched automatically (numerical, smooth) using the Leica imaging software (LAS AF-2.7.3.9723). Images were analysed on FiJi software (ImageJ). For analysis of follicles, GCs and the FDC network, tissues sections were blocked and permeabilised with 1% BSA and 2% Triton X-100 for 1hr at room temperature. Sections were stained with IgD-APC, CD21/25-biotin and chicken anti-GFP overnight at 4°C. Sections were washed thrice with PBST 0.05% Tween-20 and incubated with 1ug/mL strepadivin-AF555 and goat anti-chicken-488 (both Thermofisher) for 2hr at 4°C. Sections were washed in PBST and mounted in Hydromount (National Diagnostics) and 40X tile scanned images acquired on a Zeiss LSM-780 at Imperial College London’s FILM facility.

### Image analysis

Lymph node section fibroblast network was analysed in the parenchymal region of lymph nodes using the Mitochondrial Network Analysis (MiNA) toolset (*50*). Quantification was performed on ImageJ using the MiNA 3.0.1 macro available on https://gitgub.com/StuartLab/MiNA. Pre-processing of images was set as follows: median filter radius=5, unsharp mask filter radius=10, mask weight=0.6, CLAHE blocksize=127, histogram bins=256 and max slope=3. Data output from MiNA 3.0.1 quantified the mean branch length as the mean length of all lines used to represent the fibroblastic network structures; mean number of network branches the mean number of attached lines used to represent each structure; and the network footprint as the area or volume of the image consumed by the network signal. To quantify CD11b^+^ cell number and circularity index in the lymph node parenchyma, uniformly sized maximum projections were acquired and processed in Fiji (ImageJ) with smoothing, Gaussian blur (radius =2), and Otsu thresholding. The ‘Fill Holes’ and ‘Watershed’ functions were then applied, and particles between 20-100um^2 were analysed for cell count and circularity. For analysis of B follicles and GCs, antibody staining was thresholded and masked using a custom ImageJ macro to define individual B cell follicles (IgD) and the FDC network (CD21/35). GCs were manually defined as IgD-negative within IgD^+^ follicles and containing FDCs. The follicle and GC area and size of the corresponding FDC network were determined using the measurement tool (ImageJ). Image analysis was performed on at least 4 different lymph nodes from independent experiments.

### Serum ELISAs

ELISA plates were coated with 50 μl/well of Ovalbumin at 100 μg/ml or PBS and incubated at 4°C overnight. Plates were washed twice with PBS-Tween and then blocked for 2 hours at room temperature on a shaker with 100 ml/well with PBS containing 5% powdered milk. After washing twice with PBS-Tween, 2-fold dilutions of serum were prepared in blocking buffer at a starting dilution of 1:10, adding 100 μl/well for 2 hours at room temperature on a shaker. Wells were washed three times with PBS-Tween and 50 μl/well of biotinylated detection antibody was added (0.5 μg/ml in 1% BSA) for 1.5 hours at temperature on a shaker. Antibodies IgG1 (BD553441, clone – A85-1), IgG2a (BD553388, clone – R19-15) and IgE (BD553419, clone – R35-118). Plates were washed 5 times with PBS-Tween and 50 μl/well of Streptavidin-HRP (Sigma) was added for 30 minutes at room temperature in the dark. After washing 3 times with PBS-Tween, 100 μl/well of TMB substrate (Life Technologies) followed by 50 μl/well of 1M H2SO4 (stop substrate) were added. Optical density was read at 405 nm on a SpectraMax Plus plate reader.

### Western Blotting

Control FRCs or shRNA knockdown (PDPN-KD) cells were previously described by Acton et al. 2014(*6*). Cells were grown to ∼90% confluency in 145 mm plates and washed in ice-cold 1X tris-buffered saline pH 7.4 (TBS), scraped from dishes on ice and collected in micro-centrifuge tubes in 2 ml of ice-cold TBS. After 10 second table-top centrifugation, supernatants were removed, and cell pellets were triturated in 600 μl of ice-cold 0.1% NP-40. Protein was quantified using 5X Protein Assay Dye Reagent Concentrate (BioRad, 5000006). 6X Laemmli reducing sample buffer was added to 1X final concentration followed by boiling at 95°C for 10 min. Samples were loaded and electrophoresed using 4-20% sodium dodecyl sulphate polyacrylamide gel electrophoresis (SDS-PAGE) and transferred to PVDF membranes for 2 hrs 30 min at 65 V. Membranes were blocked in 2.5% milk and 2.5% BSA in 0.1% TBS-Tween (TBS-T), for 1hr at room temperature. Membranes were incubated with antibodies in 0.5% milk and 0.5% BSA in 0.1% TBS-T at 4°C overnight. Membranes were washed with 0.1% TBS-T followed by incubation for 2hrs with HRP-conjugated secondary antibodies in 0.5% milk and 0.5% BSA in 0.1% TBS-T. After washing with TBS-T, target protein signals were detected by ECL (GE Healthcare) and visualised using an ImageQuant (GE Lifesciences).

### Bulk RNA-seq and reads processing

RNA sequencing data from Martinez et al., 2019 are publicly available through University College London’s research data repository (doi:10.5522/04/c.4696979). Differential expression analysis was performed using R 4.0.5 and the DESeq2 package. The gene raw count table (*34*) was used to calculate differences in transcript expression between samples. As recommended in the package documentation, transcripts with fewer than 10 counts in total were removed from differential expression analysis. In all, 21285 genes were kept after prefiltering. DESeq2 was used to calculate differences in transcript expression in either PDPN-WT vs PDPN-KO samples, PDPN-WT no CLEC-2 vs CLEC-2 at 6 hours, or PDPN-WT no CLEC-2 vs CLEC-2 at 24 hours (using the contrast function). Differentially expressed genes for each comparison were defined as those with an adjusted p-value smaller than 0.05 and a fold change log2FC<1.6. (downregulated) or log_2_FC >1.6 (upregulated). The dplyr package was used for filtering the genes according to their adjusted p-values and fold change. Venn diagrams were plotted using the VennDiagram package. Heatmaps were plotted using TPM (transcript per million) values normalised per gene (Z-score) and clustered using hierarchical clustering using the pheatmap and hclust R packages. Colour palettes were generated using the RColorBrewer package. DAVID (*67*) was for functional annotation. The list of differentially expressed genes (using official gene symbols) was uploaded to the DAVID website (https://david.ncifcrf.gov/summary.jsp). The table output was then processed using a custom R script. The terms were filtered by fold discovery rate (FDR≤0.05), sorted by fold enrichment and plotted using the dplyr and ggplot2 packages. The annotated GO terms were used to identify the transcription factors in the list. Genes were mapped to GO terms using the biomaRt package for R. Transcription factors were then filtered as genes mapped with the GO:0003700 (DNA-binding transcription factor activity) term. Fold change values were plotted using the ggdotchart package. Functional network analysis was carried out using STRING. The lists of differentially expressed genes were submitted to the STRING website (string-db.org/) and analysed using the default values except for the confidence threshold and retaining physical and functional interactions. The confidence threshold was set to the most restrictive value (“very high”, 0.9). The networks were then analysed in Cytoscape (*68*) for visualisation and clustering. Nodes that did not form part of a larger network were hidden (retaining nodes with at least 3 neighbours at level 3). Clustering was performed with the MCODE algorithm (*69*) using the default parameters. Clusters were manually labelled based on the annotated gene names and descriptions. We filtered the GO terms by statistical relevance and sorted them by fold enrichment (mapped/expected).

### scRNA-seq analysis

Inguinal, axillary and brachial lymph nodes were collected from PDGFRα^mGFP^ (Control) and PDGFRα^mGFPΔPDPN^ mice (6 mice per group) and an identical number of cells was pooled from each replicate. Cells were separated by autoMACS (following manufacturers protocol) sorting into two groups: CD45^-^CD31^-^ and CD45^+^CD31^+^. Purity check of the enriched cells was performed by flow cytometry. Cells were divided into 4 groups: Control_CD45^-^CD31^-^, ΔPDPN_CD45^-^CD31^-^, Control_CD45^+^CD31^+^ and ΔPDPN_CD45^+^CD31^+^. Cells were processed through the Chromium Single-Cell 3’ v2 Library Kit (10x Genomics) at QMUL, Genome Centre. After quality control, 12825 CD45^-^ CD31^-^ cells (fibroblasts) from the Control mice (median genes/cell= 3213, median reads/cell = 43452) and 10246 CD45^-^CD31^-^ cells (fibroblasts) from the ΔPDPN mice (Median genes/cell=3567, median reads/cell = 43452) were analysed. For CD45^+^CD31^+^ enriched cells, after quality control, 6212 CD45^+^ and CD31^+^ cells from the PDGFRα^mGFP^ (Control) mice (median genes/cell= 2227, median reads/cell = 49746) and PDGFRα^mGFPΔPDPN^ (ΔPDPN) mice (Median genes/cell=2439, median reads/cell = 42869) were analysed. All scRNA-seq analysis for datasets, and subcluster analysis, was performed using the filtering, normalisation and marker identification codes previously published by Seurat et al. using Seurat (*70*–*74*). Clustering of cells was performed on merged datasets of PDGFRα^mGFP^ (Control) and PDGFRα^mGFPΔPDPN^ (ΔPDPN) cells. For cell communication stromal cells were set as the sources and immune cells as the targets. *Milo* was utilised to measured ifferential abundance analysis on KNN graphs (miloR) (*41*). Pseudotime trajectory analysis was performed on UMAP embedding using Monocle 3 v.1.0.0 package and the *Cd34*^+^ SC subcluster expressing *Pi16* neighbourhood was selected as the root (*75*). For analysis combining both CD45^-^CD31^-^ and CD45^+^CD31^+^ datasets, 6212 per condition were down-sampled and merged before analysis (Seurat code). Cell-cell communication between stromal and immune cells in PDGFRα^mGFP^ (Control) and PDGFRα^mGFPΔPDPN^ was measured using CellChat (*55*, *56*). No new codes were developed to perform analysis in this manuscript.

### Prism and statistics

Prism9 Software (GraphPad) was used to create graphs and statistical analysis. Comparisons of two data sets was performed using two-tailed Mann-Whitney tests. For serum ELISA the Wilcoxon test was used and p-values are indicated. For comparisons of multiple groups was performed using one- or two-way ANOVA with Tukey’s multiple comparison. For all tests, p<0.05 was considered significant

## Acknowledgements

We thank H. Clevers for supplying R26R-confetti mice and S. Watson and C. Buckley for providing the PDPN^fl/fl^ mouse model. We thank the CRUK Centre for Flow Cytometry TTP at UCL Cancer Institute and their staff for support and instrumentation. We thank the Genome Centre at QMUL for processing through the Chromium Single-Cell3’ v2 Library Kit (10x Genomics) and performing initial quality control and preparing gene expression libraries. We also thank Professor Sanjiv Luther and (University of Lausanne), Dr. Nagham Alouche (University of Lausanne) and Dr. Delan N. Alasaadi (UCL, LMCB) for critical discussion of data during manuscript preparation.

## Funding

This work has been supported by the European Research Council Starting Grant LNEXPANDS (S.E.A.), Cancer Research UK Careers Development Fellowship CRUK-A19763 (S.E.A.), Rosetrees Trust Foundation PGS22 100023-CD1 (S.E.A.), Cancer Research UK Senior Fellowship CRUK - RCCSCF-May22\100001 (S.E.A.) and MRC Career Development award MR/V009591/1 (AED). The LMCB was supported by Medical Research Council (MC-U12266B).

## Author contributions

S.M. and S.E.A. designed the study and wrote the manuscript. S.M. performed all *in vivo* experiments, analysed the majority of experiments and performed all scRNA-seq analysis from *in vivo* experiments. Y.H-G., J.A.C-R., V.G.M., H.L.H. and C.M.d.W. performed *in vitro* bulk-RNA-seq analysis. A.C.B. and S.E.A. designed and validated the PDGFRa-mGFP-CreERT2 mouse model. M.J. performed skeletisation image analysis. D.S. performed image analysis of CD11b cells and N.N. performed western blot analysis. I.C and A.D. performed analysis of B cell follicles and Germinal Centres.

## Competing Interests

The authors declare they have no competing interests.

## Data and Materials Availability

Bulk RNA sequencing data from Martinez et al., 2019 are publicly available through University College London’s research repository (doi:10.5522/04/c.4696979). scRNA-seq data is available through University College London’s research repository (under CC0 license).

## Supplementary Figure

**Fig. S1.**
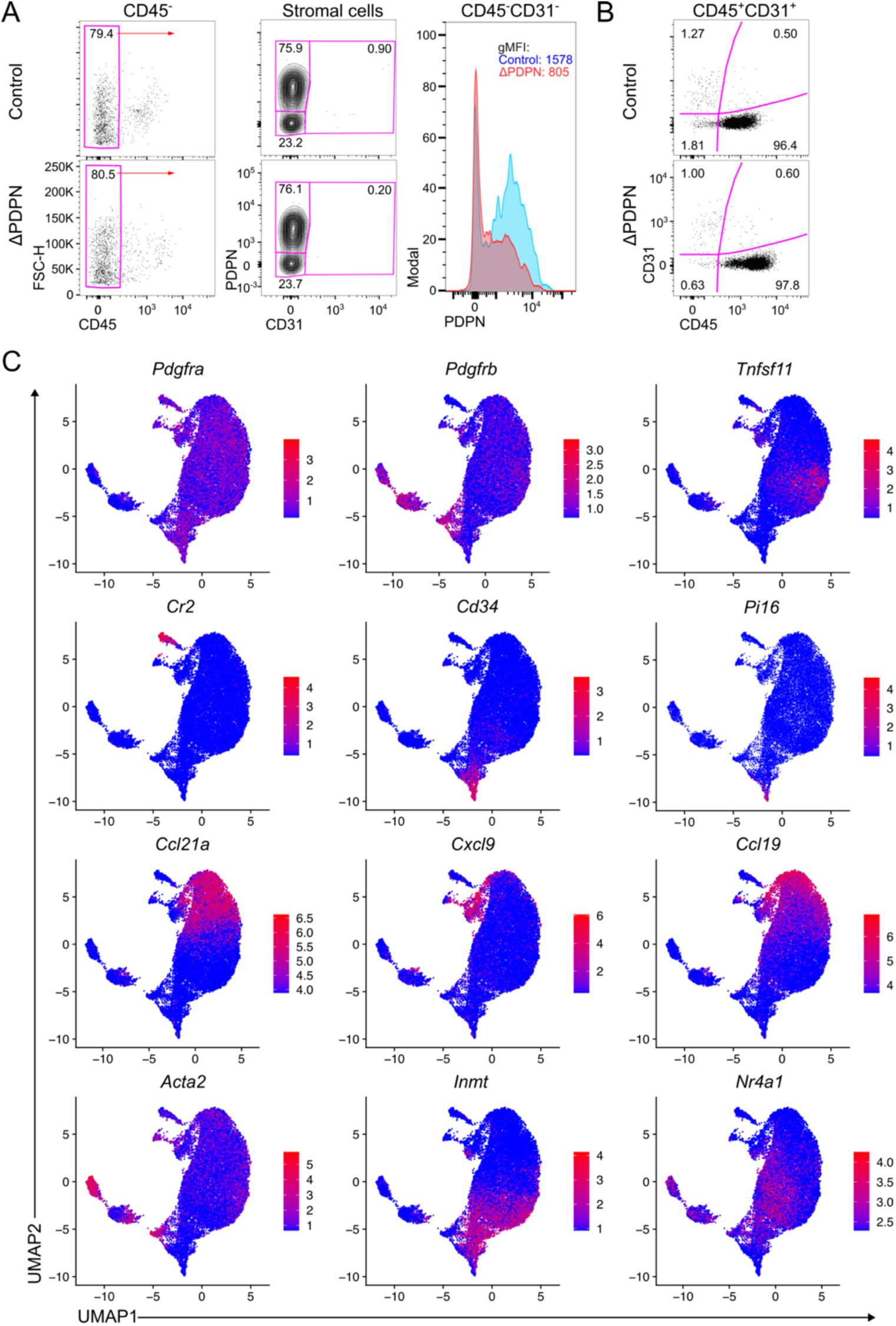
Cell purity check and gene expression on scRNA-seq analysed stromal cells. (**A**) Enriched stromal cells (CD45^-^CD31^-^) from PDGFRα^mGFP^ (control) or PDGFRα^mGFPΔPDPN^ (ΔPDPN) showing purity check (markers: CD45, CD31, PDPN) and histogram of PDPN expression with geometric mean fluorescence intensity (gMFI). (**B**) CD45^+^CD31^+^ (immune and endothelial) enriched cell purity check (markers CD45 and CD31) for PDGFRα^mGFP^ (control) or PDGFRα^mGFPΔPDPN^ (ΔPDPN) cells. (**C**) UMAP showing the expression of the primary genes used to define the stromal cell (CD45^-^ CD31^-^) clusters.

**Fig. S2.**
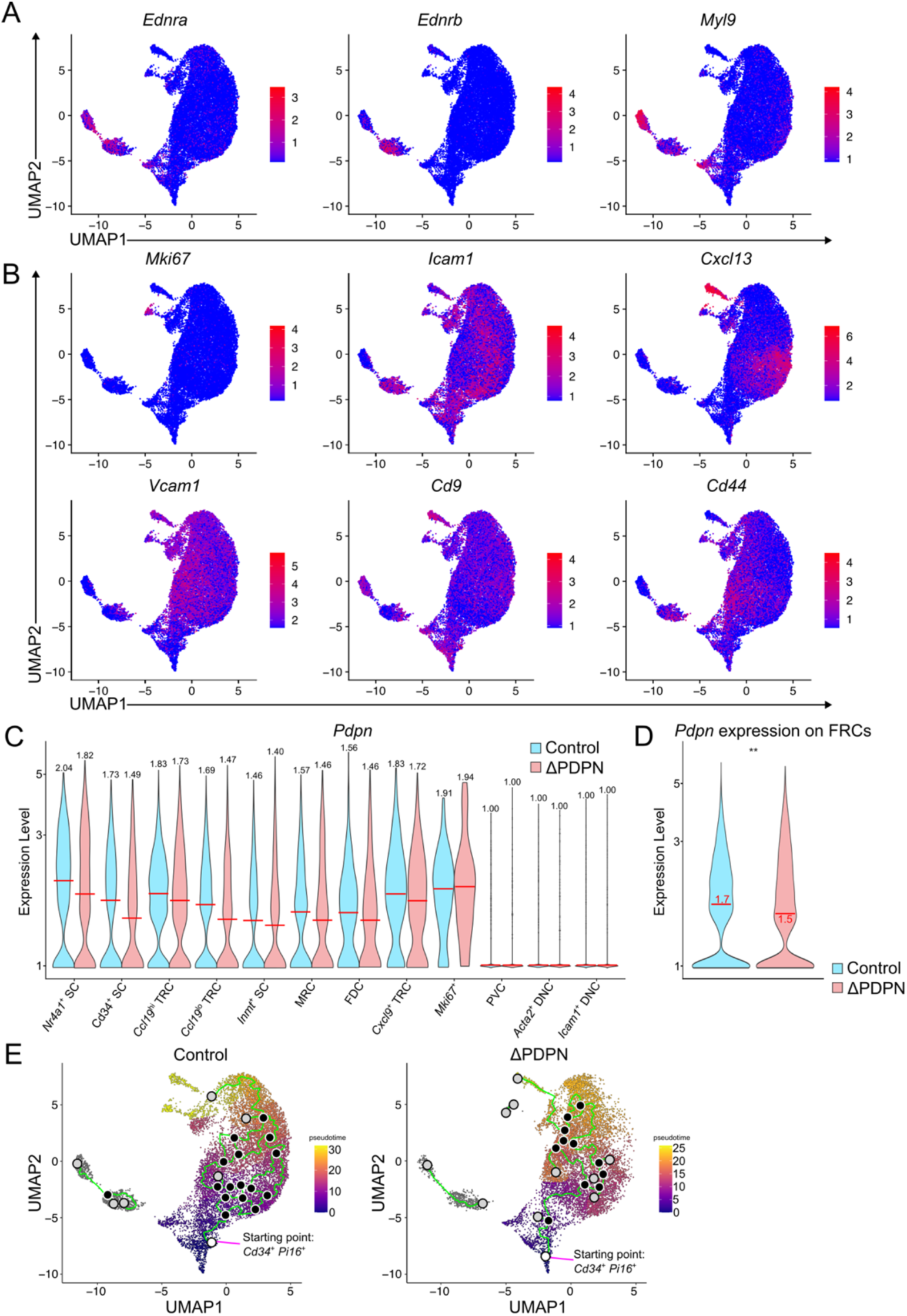
Fibroblast gene expression and trajectory analysis. (**A**) Expression on fibroblasts UMAPs for genes characterising PDGFRβ^+^ fibroblasts. (**B**) Gene expression on fibroblast UMAPs for markers defining the fibroblasts clusters. (**C**) expression of *Pdpn* across all fibroblast clusters in PDGFRα^mGFP^ (control) or PDGFRα^mGFPΔPDPN^ (ΔPDPN) cells and (**D**) on PDGFRα cells. Mann-Whitney test (two tailed), ns = no significance, **p<0.01. (**E**) Trajectory analysis (Monocle3) PDGFRα^mGFP^ (control) or PDGFRα^mGFPΔPDPN^ (ΔPDPN) with starting point at *Cd34*^+^*Pi16*^+^ stromal cells (white circle), black circles indicate branch points and light grey circle correspond to a cell fate.

**Fig. S3.**
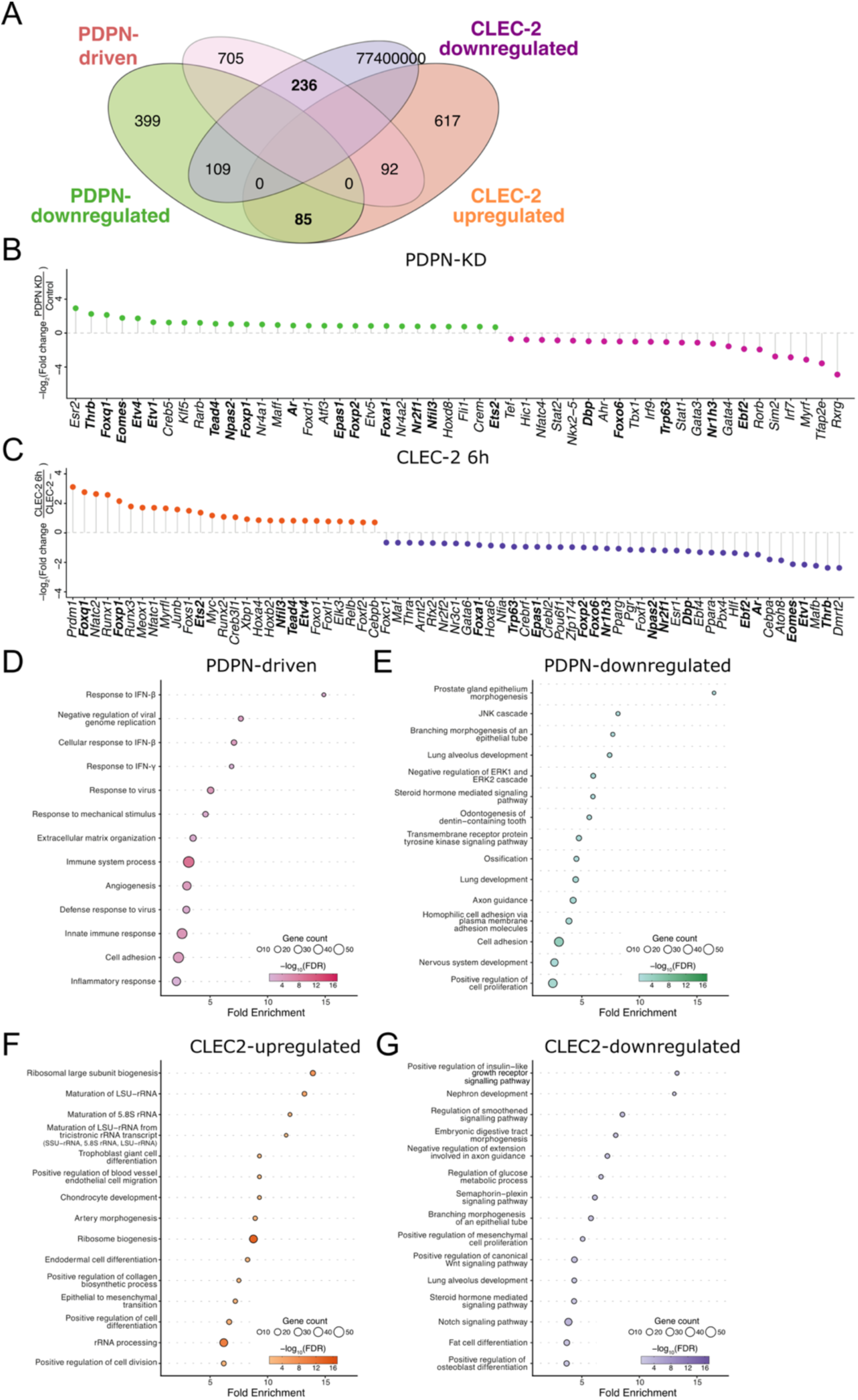
Functional annotation of the PDPN/CLEC-2 differentially expressed genes. (**A**) Venn diagram of the differentially expressed genes regulated by PDPN-KD or treatment with CLEC-2 after 6h. Transcription factors regulated by (**B**) PDPN-KD or (**C**) CLEC-2 treatment for 6 hours. (**D-G**) Gene enrichment using the GO Term database on DAVID. (**D**) PDPN downregulated (**D**) CLEC-2 upregulated (**F**) and CLEC-2 downregulated (**G**) pathways. Dot colour represents statistical significance (log2 FDR) and size is proportional to gene count.

**Fig. S4.**
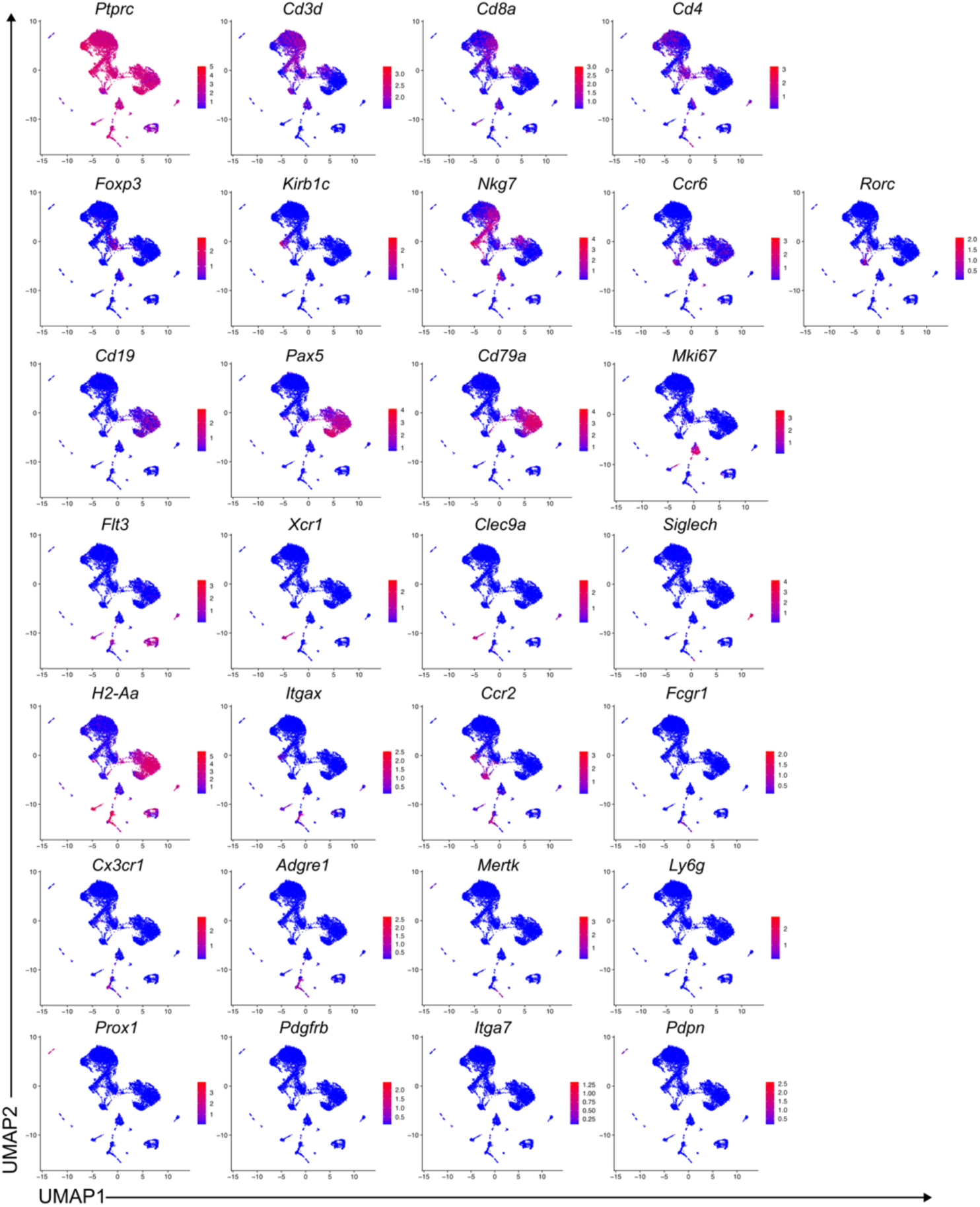
Expression of genes in CD45^+^ and CD31^+^ enriched cells. UMAP showing key genes used to define CD45^+^CD31^+^ (immune and endothelial cell) clusters.

**Fig. S5.**
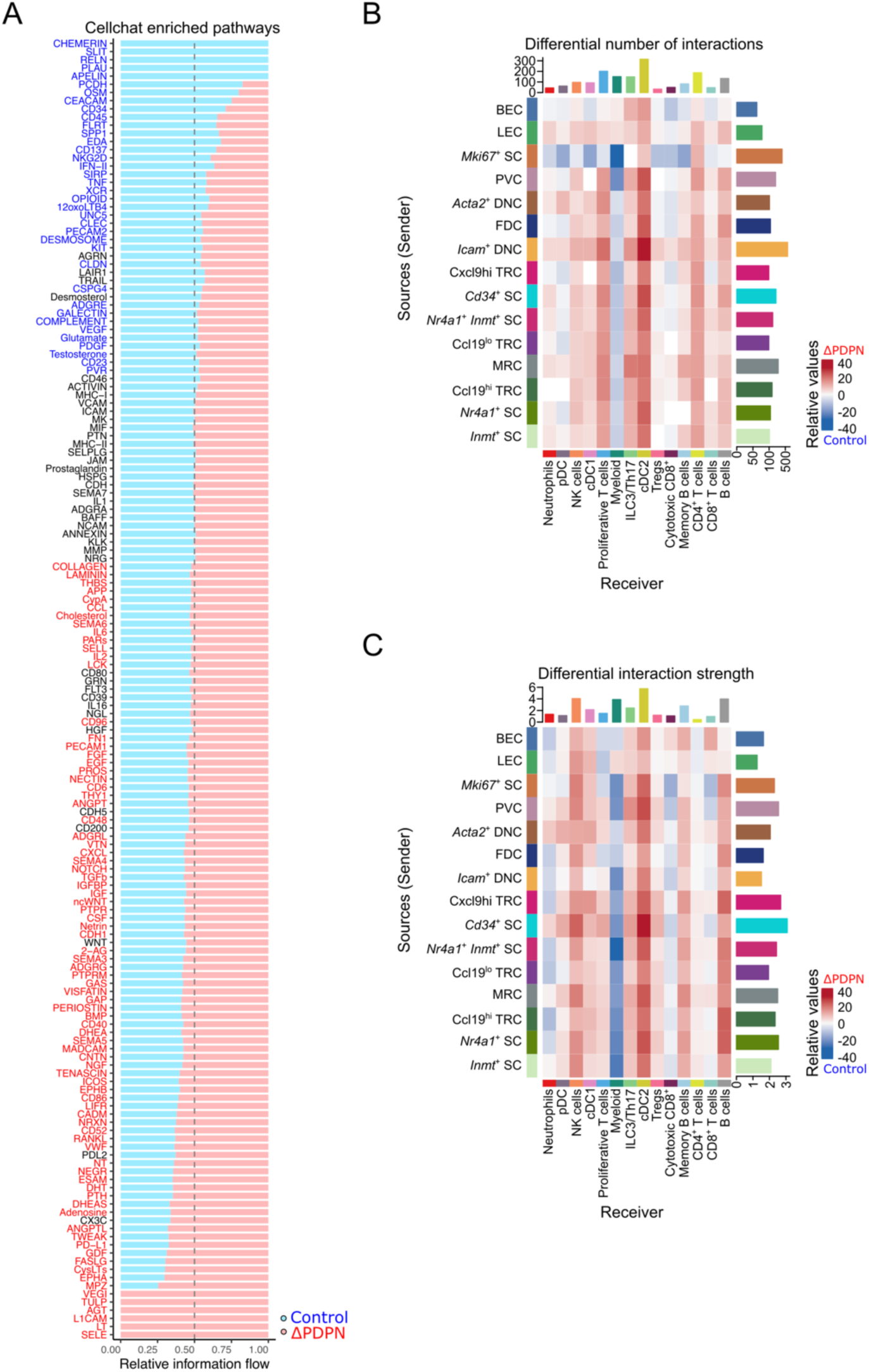
Cellhat analysis on communication between stromal and immune cells. (**A**) Cellchat enriched pathways between PDGFRα^mGFP^ (Control) or PDGFRα^mGFPΔPDPN^ (ΔPDPN) in merged CD45^-^CD31^-^ and CD45^+^CD31^+^ datasets. (**B**) Number of interactions and (**C**) interaction strength between stromal and immune cells. CellChat scores between stromal cells (source) and immune cells (receiver) where control is shown in blue and ΔPDPN in red.

**Fig. S6.**
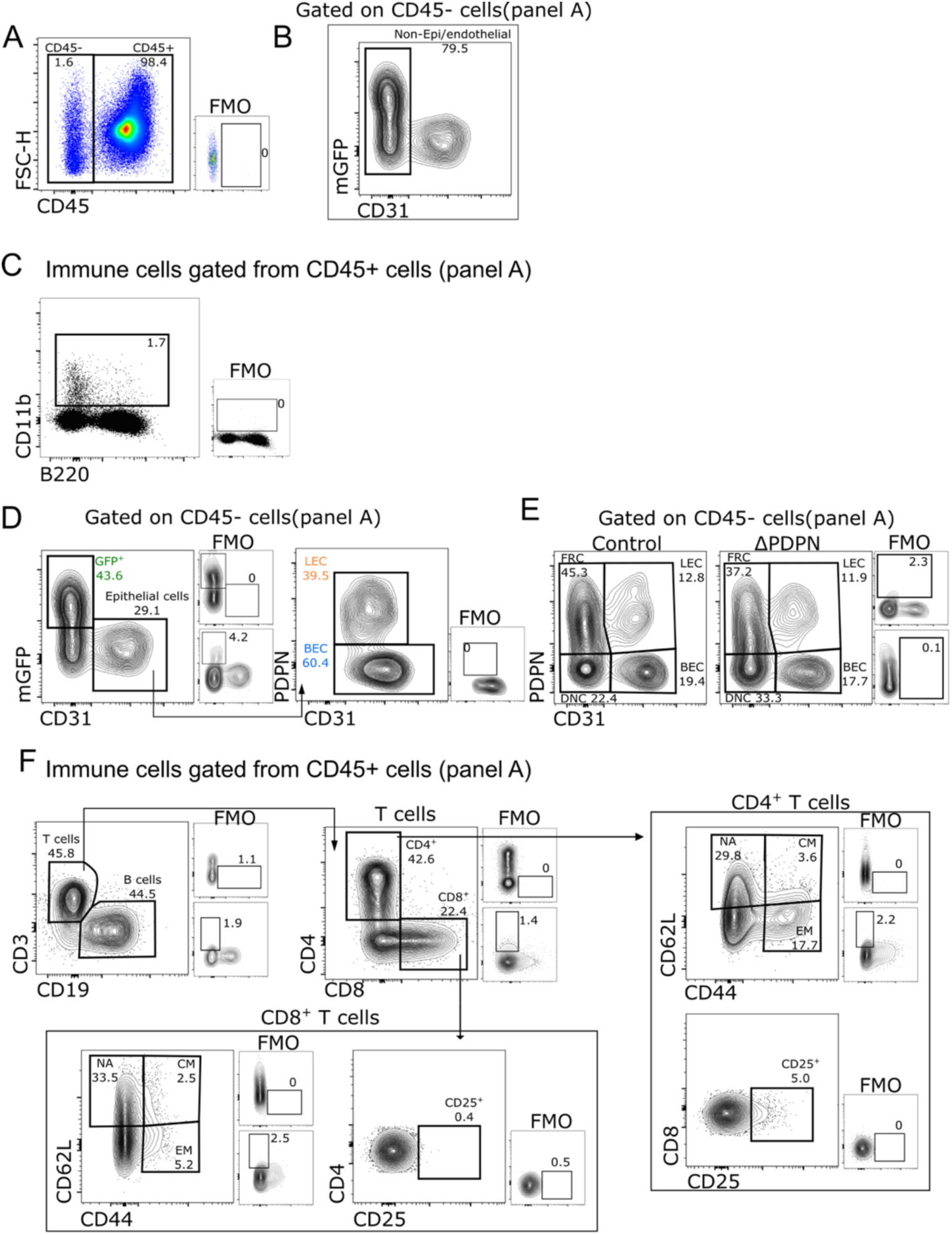
Gating strategy for stromal and adaptive cells. Flow cytometry gating used to define (**A**) CD45^-^ or CD45^+^ cells. (**B**) Gating of myeloid cells and (**D-E**) gating used to define FRCs, LECs, BECs and DNCs. (**F**) Gating strategy for defining T adaptive cell subsets.

**Fig. S7.**
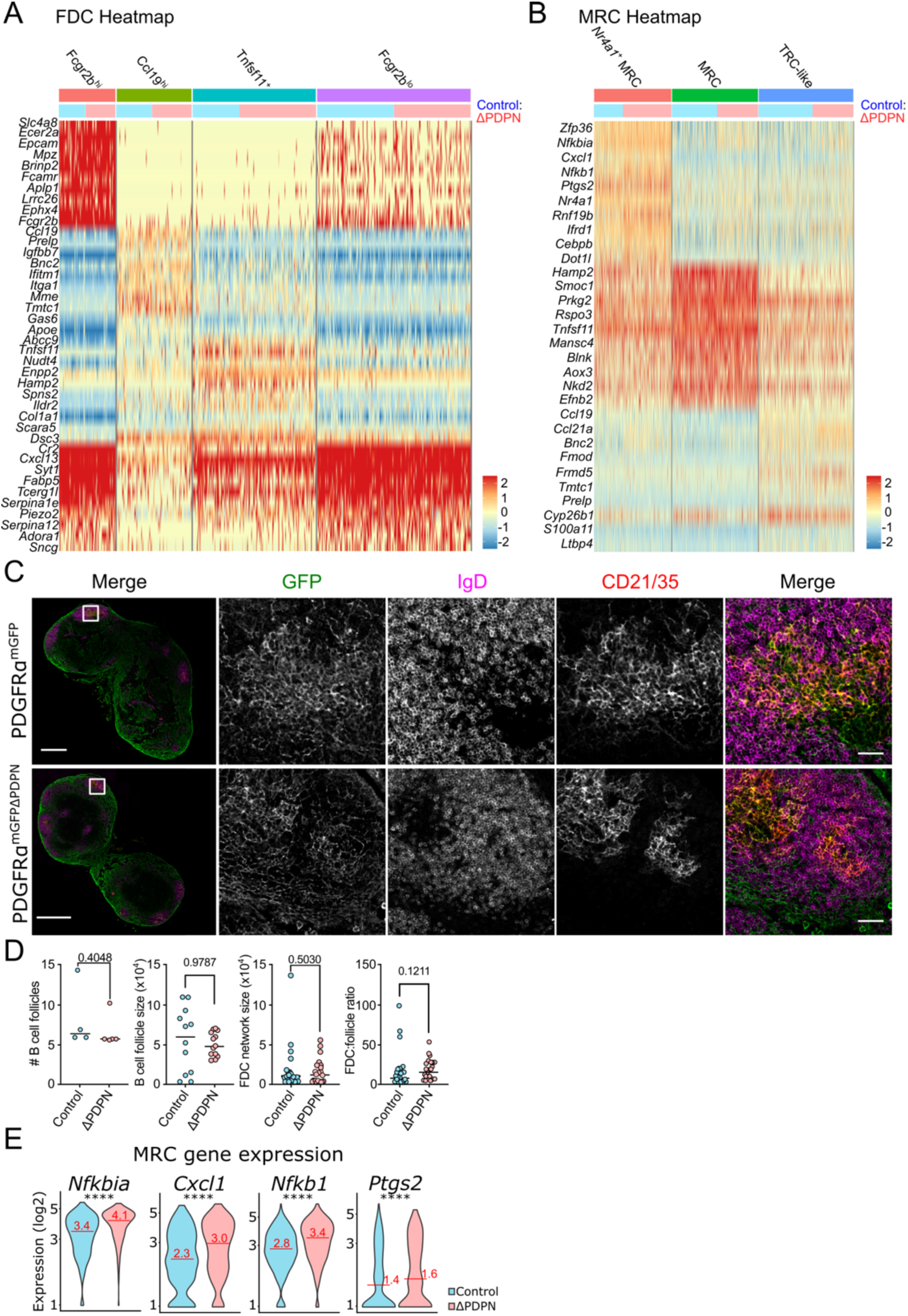
B cell follicle network organisation and FDC/MRC phenotypes altered in PDGFRα^mGFPΔPDPN^ lymph nodes. (**A**) FDC and (**B**) MRC clusters showing Heatmap showing top 10 DEG for each defined subcluster. Ratio of Control (blue) and ΔPDPN (pink) numbers (top row). (**C**) Tilescans of non-immunised PDGFRα^mGFP^ or PDGFRα^mGFPΔPDPN^ lymph nodes staining for PDGFRα (GFP), IgD (magenta), CD21/35 (red). (**D**) Analysis of number of B cell follicles, follicle size, FDC network size and FDC to follicle ratio. Mann-Whitney test (two tailed). Scale bar in tilescan is 500 μm and 50 μm in zoomed images. Images representative of 4-5 lymph nodes for each condition from 2 independent experiments. (**E**) Expression of NF-κB pathway genes enriched in MRC cluster. Mann-Whitney test (two tailed), ****p<0.0001.

**Fig. S8.**
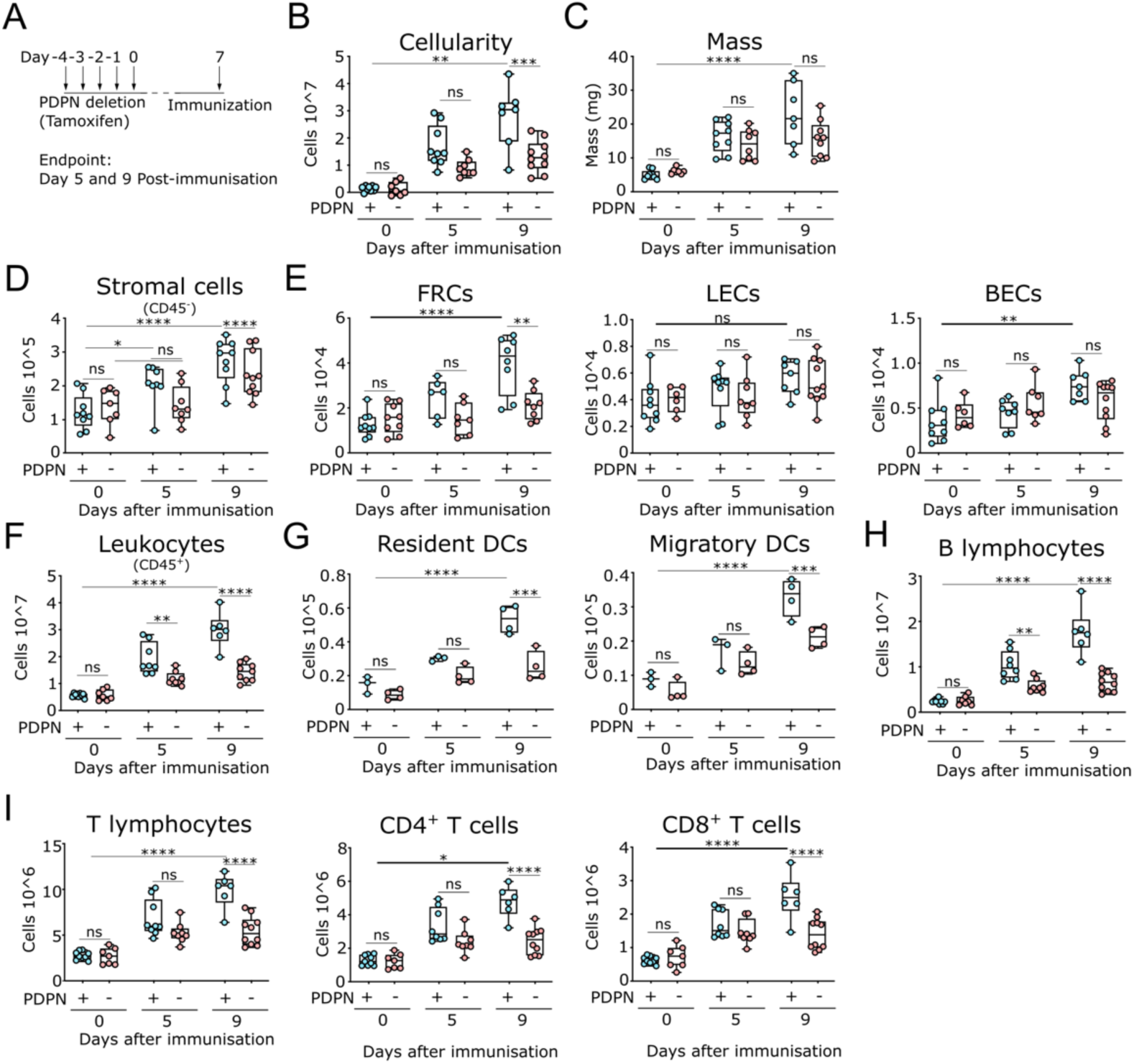
Lymph node expansion after immunisation is reduced by PDPN deletion. (**A**) Schematic of experimental design for PDGFRα^mGFP^ (Control) or PDGFRα^mGFPΔPDPN^ (ΔPDPN) mice immunisation with Incomplete Freund’s Adjuvant with ovalbumin (IFA-OVA). (**B**) Total cellularity and (**C**) lymph node mass measurement. Numbers of (**D**) stromal cells number (**E**) FRCs, LECs, BECs, (**F**) leukocytes, (**G**) Resident DCs, Migratory DCs, (**H**) B lymphocytes, (**I**) T lymphocytes, CD4^+^ T cells and CD8^+^ T cells. Flow cytometry gating found on fig. S6. N= 3-5 mice per timepoint representative of 3 independent experiments. Mann-Whitney test (two tailed), ****p<0.0001, ***p<0.001, **p<0.01, *p<0.05, ns=no significance.

## Supplementary Tables

Table S1. Gene expression for each fibroblast cluster with condition comparison.

Table S2. Differentially expressed gene for each fibroblast cluster.

Table S3. Gene expression and proportion between conditions in fibroblast clusters.

Table S4. *Nr4a1*^+^ subcluster differentially expressed genes.

Table S5. *Ccl19*^hi^ subcluster differentially expressed genes.

Table S6. Gene overlap between *in vivo* scRNA-seq and *in vitro* bulk RNA-seq.

Table S7. Comparison of gene expression and DESeq *in vitro*.

Table S8. STRING analysis *in vitro*.

Table S9. Differentially expressed genes in CD45^+^ and CD31^+^ cells.

Table S10. Myeloid and macrophage subcluster differentially expressed genes.

Table S11. Dendritic cell subcluster differentially expressed genes.

Table S12. B cell subcluster differentially expressed genes.

Table S13. FDC subcluster differentially expressed genes.

Table S14. MRC subcluster differentially expressed genes.

Table S15. Materials and consumables.

## Notes

### Competing Interest Statement

The authors have declared no competing interest.

### Summary of Updates

The rewrite of this paper reflects the addition of extensive RNAseq analysis, which extends some of our previous conclusions. The result is several new figures, and some original figures are removed or moved to supplementary data.

